# Rapid elicitation of a new class of neutralizing N332-glycan independent V3-glycan antibodies against HIV-1 in nonhuman primates

**DOI:** 10.1101/2025.09.04.674340

**Authors:** Ignacio Relano-Rodriguez, Jianqiu Du, Zi Jie Lin, Margaret Kerwin, Marta Tarquis-Medina, Eduardo Urbano, Jiayan Cui, Meagan Watkins, Peng Zhao, Rumi Habib, Sukanya Ghosh, Joyce Park, Caroline Boroughs, Agnes A. Walsh, Mariane B. Melo, George M. Shaw, Beatrice H. Hahn, Darrell J. Irvine, Lance Wells, David B. Weiner, Daniel W. Kulp, Ronald S. Veazey, Jesper Pallesen, Amelia Escolano

**Affiliations:** Vaccine and Immune Therapy Center, The Wistar Institute, Philadelphia, PA 19104, USA; Department of Molecular and Cellular Biochemistry, Indiana University Bloomington, IN 47405, USA; Department of Chemistry, Indiana University Bloomington, IN 47405, USA; Department of Bioengineering, University of Pennsylvania, Philadelphia, PA, USA; Division of Comparative Pathology, Tulane National Primate Research Center, Covington, LA 70433 USA; Complex Carbohydrate Research Center, University of Georgia, 315 Riverbend Road, Athens, GA 30602, USA; Department of Medicine, University of Pennsylvania, Philadelphia, PA, USA; Department of Microbiology, University of Pennsylvania, Philadelphia, PA, USA; Department of Immunology and Microbiology, Scripps Research Institute, La Jolla, CA, USA; Howard Hughes Medical Institute, Chevy Chase, MD, USA

## Abstract

Sequential immunization is a promising approach to elicit broadly neutralizing antibodies (bNAbs) against the HIV-1 Envelope (Env). However, available protocols are inefficient and involve multiple immunizations over long periods of time. Here, we present WIN332, a new engineered Env-immunogen that induces a new class of neutralizing N332-glycan-independent antibodies to the conserved V3-glycan epitope of Env after a single bolus immunization in nonhuman primates. WIN332 binds to precursors of canonical human N332-glycan-dependent (Type-I) V3-glycan bNAbs but also of a first-of-its-class N332-glycan-independent (Type-II) V3-glycan bNAb. A single immunization elicits neutralizing serum and monoclonal antibodies that are boosted and affinity matured with a heterologous immunogen. EMPEM analysis of serum antibodies, antibody cloning and cryo-EM analysis reveal that WIN332 elicits N332-glycan-independent antibodies with remarkable sequence and binding similarities with the most potent human type-I and type-II V3-glycan bNAbs. Thus, WIN332 is a promising vaccine candidate to streamline V3-glycan bNAb elicitation.

## Main

Sequential immunization using a series of engineered and native-like HIV-1 Envelope (Env) proteins is a promising approach to eliciting broadly neutralizing antibodies (bNAbs) against HIV-1^1^. Sequential immunization aims to activate naïve B cells expressing bNAb precursor antibodies and subsequently induce affinity maturation to increase antibody neutralization potency and breadth. To date, only two sequential immunization protocols designed to induce antibodies against the fusion peptide and CD4 binding site of Env have consistently induced bNAbs in nonhuman primates (NHPs)^2,3^. However, the elicited responses showed limited potency and breadth and required multiple immunizations over long periods of time.

Different bNAb lineages have different constraints for maturation. The activation of “premium” antibody lineages that require less maturation to acquire neutralization potency and breadth would simplify sequential immunization protocols. In this regard, the conserved V3-glycan epitope of Env is the most common specificity for broadly neutralizing activity in studies of cohorts with bNAb responses^4,5^. The V3-glycan epitope comprises a peptide motif, GDIR, and several surrounding glycans including those at positions N133, N137, N156, N301 and N332^6^. Interaction with the N332-glycan is a defining signature of V3-glycan bNAbs^7–9^ and a requirement for V3-glycan bNAbs to neutralize HIV-1, thus all currently available immunogens to elicit V3-glycan bNAbs are designed to retain this glycan^10–14^. However, we hypothesize that N332-glycan dependency develops during bNAb maturation and that unmutated precursors and early intermediates of V3-glycan bNAbs may not require this glycan for binding and activation. We further hypothesize that N332-glycan-independent antibody lineages can mature into N332-dependent (type-I) and N332-glycan-independent (type-II) V3-glycan bNAbs. Notably, a new human V3-glycan bNAb, EPTC112 has been recently isolated from an HIV infected individual^15^. EPTC112 targets the V3-glycan epitope of Env, however, the cryo-EM structure shows no contacts with the N332-glycan. This bNAb is therefore evidence that N332-glycan independent bNAbs can develop in humans.

Here, we present WIN332, a N332-glycan-deficient priming immunogen rationally designed to broadly activate precursors of both types of V3-glycan bNAbs, N332-glycan dependent (type-I) and N332-glycan independent (type-II). WIN332 prime elicits antibodies that closely resemble human type-I and type-II V3-glycan bNAbs in NHPs. Remarkably, one single bolus immunization with WIN332 elicits serum and monoclonal antibodies that neutralize wild-type fully glycosylated viruses and are further boosted by a heterologous boosting immunogen. WIN332 accelerates the development of neutralization by activating a new class of “premium” N332-glycan independent antibody lineages and arises as a promising vaccine candidate to simplify sequential immunization protocols against HIV-1.

## Results

### WIN332, an HIV-1 Env immunogen for broad elicitation of V3-glycan antibodies

WIN332 is an engineered native-like Env SOSIP trimer derived from the Env protein of the clade A/E HIV-1 strain BG505^16^ (Fig.1a). To design WIN332, we reevaluated the canonical definition of the V3-glycan epitope. We hypothesized that antibody recognition of the V3-glycan epitope may not always require interaction with the N332-glycan and that N332-glycan-independent antibody lineages can be activated and matured through immunization. Supporting this hypothesis, a study has recently reported a human N332-glycan-independent bNAb, EPTC112^15^, which confirms that this class of antibody is elicited in humans. Thus, here we considered two subtypes of V3-glycan antibody lineages: Type-I, N332-glycan dependent^7,8,17–19^ and Type-II, N332-glycan independent^15^ . To test our hypothesis, we designed WIN332, a derivative of the previously described RC1 priming immunogen^10^ that lacks the signature Potential N-linked glycosylation site (PNGS) at position N332 in addition to those previously removed in RC1 (N133, N137 and N156) (Fig. 1a, Extended Data Fig.1a and Extended Data Table 1).

**Figure 1:**
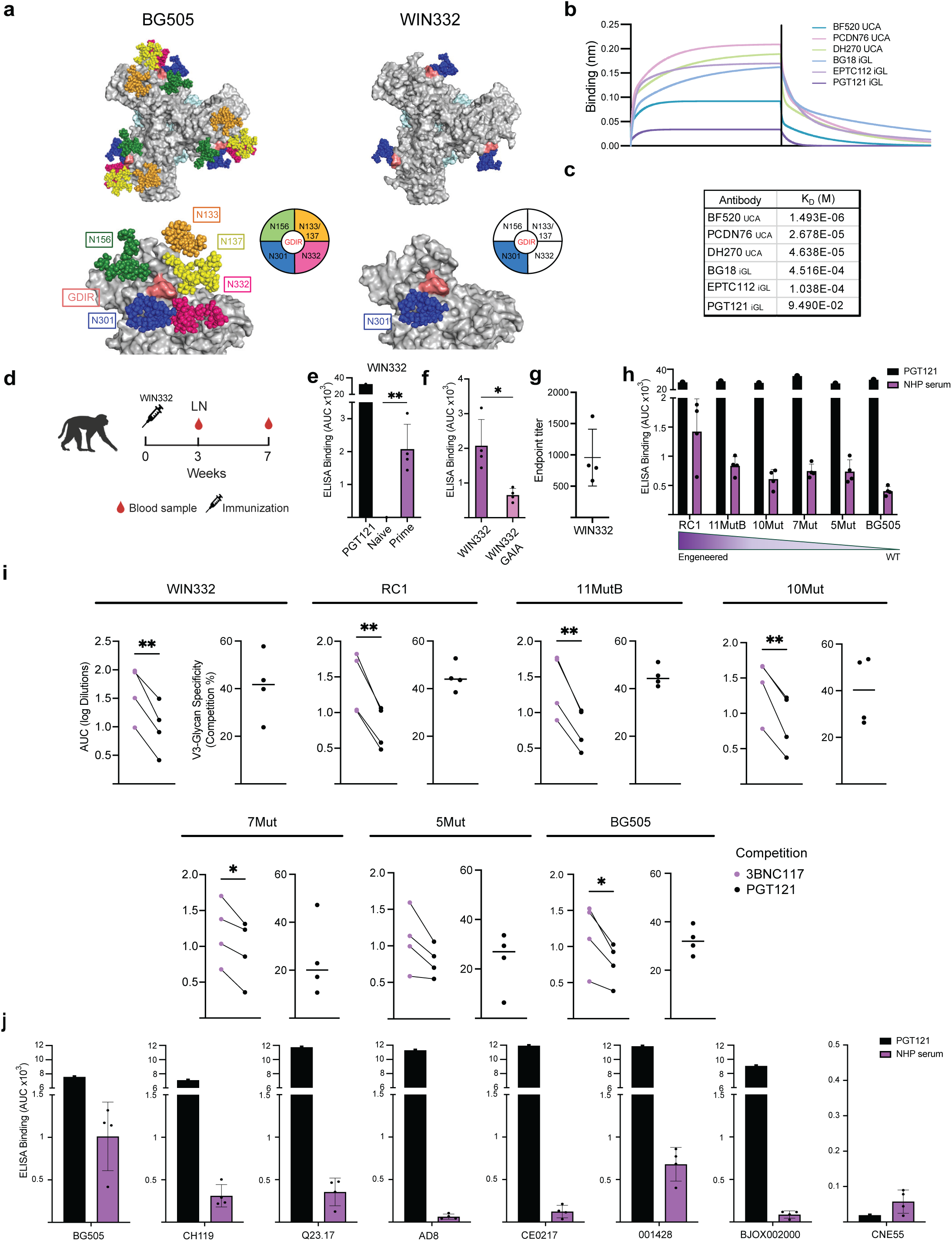
WIN332 binds to precursors of human type-I and type-II V3-glycan bNAbs and induces V3-glycan serologic responses in NHPs. **a.** Top images show the three V3-glycan epitopes of the native-like BG505 (left) and WIN332 (right) Env trimers on a top view of the trimers. Bottom images show closer views of one of the V3-glycan epitopes of BG505 (left) and WIN332 (right). Env gp120 is shown in gray, glycans are represented with colored spheres, GDIR is shown in salmon. Pie charts including colored slices are used for schematic representation of the V3-glycan epitopes. White slices indicate absent glycans. **b.** Representative BLI (OCTET) curves showing binding of WIN332 to the indicated iGL and UCA human bNAb precursors. Fabs were tested at 20 µM, 10 µM, 5 µM and 2.5 µM **c.** Table shows the calculated K_D_ for the interaction of WIN332 with the human bNAb precursors in b. **d.** Immunization protocol in NHPs. LN: Lymph node biopsy **e.** ELISA results showing binding of the serum collected from NHPs prior to immunization (Naïve) and 3 weeks after one immunization with WIN332 (Prime) to WIN332. The human V3-glycan bNAb, PGT121 was used as a positive control at 5µg/ml. **f.** ELISA results showing the binding of the serum from WIN332-primed NHPs to WIN332 and a version of WIN332 with GAIA mutations in the GDIR motif (WIN332 GAIA) **g.** Endpoint titers for ELISA in e. **h.** ELISA results showing the binding of the serum from WIN332-primed NHPs to Env proteins with serially more native and glycosylated V3-glycan epitopes. **i**. Results of competition ELISAs using the serum from WIN332-immunized NHPs on the indicated Env proteins. Human bNAbs PGT121 (V3-glycan epitope) and 3BNC117 (CD4bs) were used as competing antibodies. **j.** ELISA results showing binding of the macaque serum to different native-like Env trimers of different clades. Data were analyzed using Student’s t-test and considered statistically significant at *P ≤ 0.05 and **P ≤ 0.01.

WIN332 elutes from size exclusion chromatography columns at the expected fractions for gp140 trimers (Extended Data Fig.1b) and shows the expected propeller shape by negative staining electron microscopy (nsEM) (Extended Data Fig. 1c). Glycan site occupancy for each PNGS was determined by mass spectrometry confirming the absence of glycosylation at N133, N137, N156 and N332 and the presence of glycosylation at N301 (Extended Data Fig. 1a). WIN332 shows high binding to human bNAbs targeting the CD4 binding site (CD4bs) (VRC01), V1V2 (PGDM1400 and PGT145), V3-glycan (PGT121 and BG18) and the gp120-gp41 interface (PGT151) epitopes and undetectable or low binding to non-neutralizing antibodies including 3074, A32, F105, F425, 4025 and 17b (Extended Data Fig. 1d).

We conclude that WIN332 is a well-formed SOSIP trimer that retains the relevant antigenic properties of the HIV-1 Env.

### WIN332 binds to unmutated precursors of type-I and type-II V3-glycan bNAbs

To evaluate whether WIN332 could activate precursors of known human V3-glycan bNAbs and initiate bNAb maturation, we interrogated the binding of WIN332 to inferred germline (iGL) precursors and unmutated common ancestors (UCAs) of human type-I and type-II V3-glycan bNAbs by biolayer interferometry (Fig. 1b). Interestingly, despite the absence of the N332-glycan, WIN332 showed quantifiable affinity for the precursors of human type-I bNAbs PGT121 (*K*_D_=9.4 x 10^-2^ M), BG18 (*K*_D_=4.5 x 10^-4^ M), DH270 (*K*_D_=4.6 x 10^-5^ M), PCDN76 (*K*_D_=2.7 x 10^-5^ M) and BF520 (*K*_D_=1.5 x 10^-6^ M). Moreover, WIN332 showed quantifiable affinity for the precursor of the type-II EPTC112 (*K*_D_=1 x 10^-4^ M) bNAb (Fig.1c).

We demonstrate that WIN332 binds to precursors of V3-glycan bNAbs from both N332-glycan dependent (type-I) and independent (type-II) subtypes, suggesting that WIN332 could activate these bNAb precursors *in vivo*.

### WIN332 elicits N332-glycan-independent V3-glycan antibodies in NHPs

To characterize the ability of WIN332 to broadly elicit V3-glycan antibodies, we immunized rhesus macaques with one subcutaneous bolus injection of WIN332 in SMNP adjuvant^20^ distributed between the four limbs. Three and seven weeks after immunization, blood and draining lymph node (dLN) biopsies were collected (Fig. 1d). ELISA analysis of the serum from naïve and immunized macaques revealed a robust antibody response to WIN332 in the immunized macaques (average endpoint titer: 956) (Fig. 1e-g). The serum from WIN332-primed NHPs showed strongly reduced binding to a version of WIN332 with a mutated GDIR (WIN332 GAIA), indicating that the serum antibodies require an intact GDIR motif for binding to WIN332 (Fig, 1f and Extended Fig. 1e). The week 3 serum showed comparable binding to WIN332 and RC1, indicating that the serum antibodies could accommodate the N332 glycan. In addition, the serum showed gradually reduced binding to WIN332 variants with serially more native and glycosylated V3-glycan epitopes, including low but detectable binding to the fully glycosylated BG505 SOSIP^1,21^ (Engineering: WIN332>RC1>11MutB>10Mut>7Mut>5Mut>BG505) (Fig. 1h) (Extended Data Table 1).

To map the serologic response, we performed competition ELISAs on WIN332 using the V3-glycan bNAb, PGT121 and the CD4bs bNAb, 3BNC117 as competing antibodies (Fig.1i). Binding of the serum to WIN332 was significantly reduced (41.2%) in the presence of PGT121, suggesting that a fraction of the serum antibodies targeted the V3-glycan epitope. Competition by PGT121 was also detectable on the serially more native RC1, 11MutB, 10Mut, 7Mut and native BG505 Env proteins (Fig.1i). Remarkably, the serum from week 7 bound to a panel of native multi-clade trimers, including BG505 T332N, CH119, Q23.17, AD8, CNE55, 001428, CE0217, and BJOX002000 (Fig. 1j).

To further confirm the presence of V3-glycan antibodies in the serum of WIN332-immunized macaques, we performed electron microscopy polyclonal epitope mapping (EMPEM). EMPEM analysis using WIN332 (Fig. 2a and Extended Data Fig. 2a,b) and RC1 (Fig.2b) revealed the presence of antibodies targeting the V3-glycan epitope in the serum of the three macaques independently analyzed. Overlapping the electron density of WIN332-Fab and RC1-Fab complexes with the WIN332 and RC1 structures respectively^10^ confirmed V3-glycan specificity (Fig. 2c,d). EMPEM analysis using RC1 confirmed the presence of V3-glycan antibodies that could accommodate the N332 glycan (Fig. 2b, d, e-g). Comparison of the RC1-Fab and WIN332-Fab electron densities with published structures of Env-bound BG18, PGT121 and EPTC112^12,15,22^ showed structural alignment with all three antibodies (Fig. 2e-g and Extended Data Fig. 2c,d). In addition to V3-glycan antibodies, we detected Fabs binding the base of the Env trimer and the gp120-gp41 interface (Fig. 2a,b).

**Figure 2:**
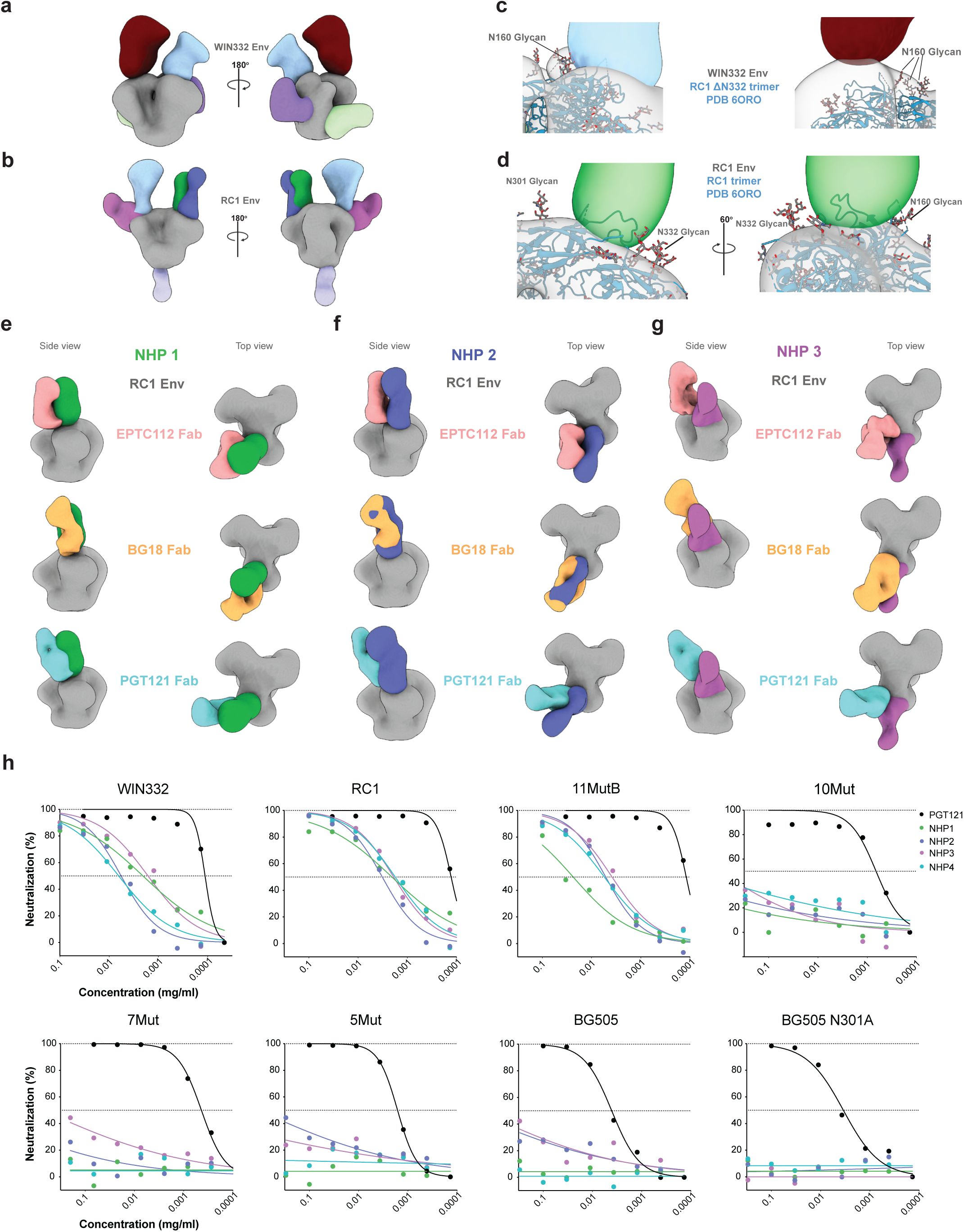
WIN332 elicits serum antibodies that target the V3-glycan epitope and neutralize the autologous fully glycosylated virus. **a, b.** EMPEM results showing the binding of Fabs (colored) purified from the serum of WIN332-primed NHP 1 and 3 to WIN332 (gray) (a) and NHP 1, 2 and 3 to RC1 (gray) (b). **c.** Close-up views highlighting the interactions of the Fabs from NHP 1 and 3 (blue and red) with the N160 glycan of WIN332. WIN332 Env density is shown in gray with the model of RC1 (PDB 6ORO) with N332 glycan removed fit in (RC1 trimer 1′N332 glycan). **d.** Densities of NHP 1 Fab (green) bound to RC1 (gray) with RC1 model fit in (PDB 6ORO). Close-up views highlighting accommodation of N332 and N301 glycans on RC1. **e-g.** Comparison of Fab densities from NHP 1 (green), NHP 2 (dark blue), and NHP 3 (purple) with EPTC112 (pink), BG18 (orange), and PGT121 (cyan) Fab densities. **h.** Representative results of TZM-bl neutralization assays using purified immunoglobulins from the serum of WIN332-primed NHPs against pseudoviruses carrying gradually more native looking Env proteins. 5MUT and BG505 (BG505 T332N) are fully glycosylated pseudoviruses. BG505 N301A is a modified version of the BG505 pseudovirus with a mutation that knocks out the V3-glycan epitope by removing the N301 glycan.

We demonstrate that one immunization with WIN332 elicits serologic antibodies targeting the V3-glycan epitope in a N332-glycan-independent manner which cross-react with sequentially more native-like Envs in NHPs.

### WIN332 elicits neutralizing serologic responses in NHPs

To evaluate the capacity of WIN332-elicited serum antibodies to neutralize HIV-1, we purified immunoglobulins from the serum from week 7 and tested them in TZM-bl assays against a panel of pseudoviruses including WIN332, RC1, 11MutB, 10Mut, 7Mut, 5Mut, fully glycosylated BG505 (BG505 T332N), a V3-glycan epitope knockout (BG505 N301A) and the Murine Leukemia Virus

(MLV) as a control (Fig. 2h and Extended Data Fig. 2e). Strong neutralization activity was observed against WIN332 and RC1 and gradually reduced activity against 11MutB, 10Mut, 7Mut and 5Mut. Remarkably, the serum showed low but detectable and reproducible neutralization activity against the autologous fully glycosylated BG505 pseudovirus (Fig. 2h). No detectable activity was observed against the V3-glycan knockout pseudovirus (BG505 N301A) and MLV control, suggesting that the serum neutralizing response was entirely immunofocused to the V3-glycan epitope (Fig. 2h and Extended Data Fig. 2e).

We conclude that one immunization with WIN332 elicits serum N332-glycan-independent V3-glycan antibodies that neutralize autologous BG505 pseudovirus in NHPs.

### WIN332 elicits N332-glycan independent antibodies that resemble type-I and type-II human V3-glycan bNAbs in NHPs

To further characterize the V3-glycan antibodies elicited by WIN332 in NHPs, we isolated single WIN332-specific or BG505-specific germinal center (GC) B-cells from dLNs collected 3 weeks after WIN332 immunization and cloned their immunoglobulin (Ig) genes (Fig 3a and Extended Data Fig. 3a). Analysis of the Ig genes revealed extensive B-cell clonal expansion (Fig. 3b ). The average number of somatic hypermutations (SHM) was 5.7 nt for VH, 2.8 for VK and 6.1 nt for VL genes (Fig. 3c). The average CDRH3 length was 14.4 amino acids (aa) ranging from 7 to 26 aa (Fig. 3d).

**Figure 3:**
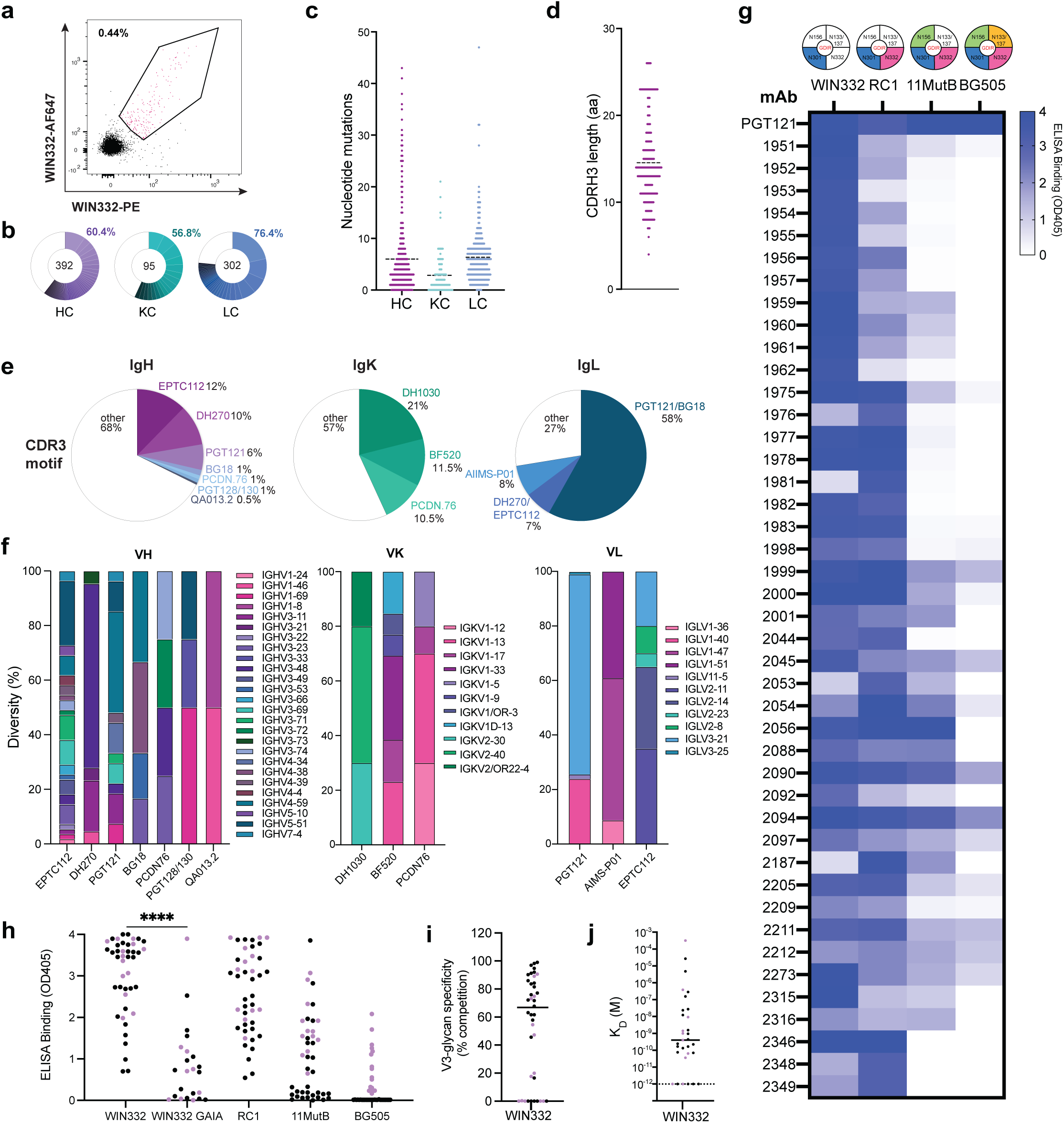
WIN332 elicits monoclonal antibodies that have sequence motifs of human type-I and type-II V3-glycan bNAbs and bind to native fully glycosylated Env. **a.** Representative flow cytometry plot showing WIN332-binding B cells in the dLNs of WIN332-primed NHPs. **b.** Pie charts show clonal expansion of WIN332-specific B cells isolated 3 weeks after WIN332 prime from NHP 2 and NHP 3. Heavy chains (HC), kappa light chains (KC) and lambda light chains (LC) are shown in purple, green and blue respectively. White slices correspond to singlet sequences. The number in the middle of the pie chart indicates the number of sequences analyzed. **c.** Nucleotide somatic hypermutations in HC, KC and LC sequences from b. **d.** CDRH3 length of sequences from b. **e.** Pie charts show the frequency of sequences with CDR3 motifs of different human type-I and type-II V3-glycan bNAbs. Data for HC, KC and LC are shown in purple, green and blue respectively. **f.** Graphs show the diversity and frequency of genes used by sequences from e. **g.** Table summarizes results of screening ELISAs to determine the binding of 43 WIN332-binding mAbs to the gradually more glycosylated Envs RC1, 11MutB and native BG505. Darker blue indicates higher ELISA binding. PGT121 was used as a positive control. **h.** Graph summarizes the ELISA binding in g, including the binding to WIN332 GAIA. BG505-binding antibodies are marked in pink in h-j. **i.** V3-glycan specificity of antibodies in g as determined by competition ELISAs on WIN332 using PGT121 (V3-glycan bNAb) and 3BNC117 (CD4bs bNAb) as competing antibodies. **j.** Affinities of antibodies in g for WIN332.

73% of the lambda light chains (IgL) presented CDRL3 motifs previously identified in human type-I and type-II V3-glycan bNAbs. These motifs included the “QxDSS” motif of type-I PGT121/10-1074/BG18 bNAbs^7,19^ and of previously reported NHP V3-glycan antibodies^10^ and the “SYxG” motif of the type-II EPTC112^15^ bNAb (Fig. 3e). 43% of the kappa light chains (IgK) presented CDRL3 motifs of human bNAbs including DH1030, BF520 and PCDN.76 ^23,24^. CDRH3 analysis showed that 12% of the 392 sequences analyzed had the key “SxW” motif of EPTC112 and 10% had the “SxYY” motif of DH270. In addition, we identified CDRH3s with the “RxY” motif of PGT121, the FGVV motif of BG18 and the “WSG” motif of AIIMS-P01 and PCDN76 ^7,9,12,15,17,25,26 24^ (Fig. 3e and Extended Data Fig. 3b). We conclude that WIN332 elicits antibodies that resemble human type-I and type-II V3-glycan bNAbs in NHPs.

### Macaque monoclonal antibodies elicited by WIN332

We produced a selection of 43 mAbs that had signature motifs of human V3-glycan bNAbs (Extended Data Table 2). To characterize the binding of these mAbs, we performed a battery of ELISAs against different engineered and native Envs (Fig. 3g,h). The ELISA results showed that 43/43 antibodies bound WIN332, 40/43 WIN332-binding antibodies bound RC1, which reintroduces the N332-glycan and 32/40 bound to 11MutB which reintroduces the N156-glycan. Remarkably, 13 mAbs bound to the fully glycosylated BG505 SOSIP trimer (Fig. 3g,h). We performed competition ELISAs on WIN332 using the V3-glycan bNAb, PGT121 and the CD4bs bNAb, 3BNC117 as competing antibodies. PGT121 competed the binding of 30/43 mAbs with an average competition of 74.8% and ranging from 50 to 90% (Fig. 3i). Moreover, mAbs showed significantly reduced binding to WIN332 with a GDIR motif mutated to GAIA (WIN332 GAIA) (Fig. 3h) further supporting V3-glycan epitope specificity. The affinity of the 43 WIN332-binding mAbs for WIN332 had an average K_D_ of 10^-9^ M (Fig. 3j)

We conclude that WIN332 elicits mAbs that cross-react with fully glycosylated BG505 trimer and are competed by the V3-glycan bNAb PGT121.

### WIN332 elicits neutralizing BG18-like antibodies in NHPs

We selected two NHP mAbs for further characterization. Ab1999 was selected for its high similarity with the human type-I V3-glycan bNAb, BG18. The HC of Ab1999 has 76.7% aa identity with the HC of iGL BG18 and 60.4% with the mature BG18 HC (Fig. 4a and Extended Data Fig. 4a). Remarkably, Ab1999 has the defining “FGVV” CDRH3 motif of BG18-like antibodies, which is involved in the binding and neutralization activity of BG18^7,27^. Ab1999 uses macaque VH4-106, DH3-3, JH5-1, VK1-74 and JK2*01 genes and has a 22 aa long CDRH3, only one aa shorter than BG18 (Fig. 4a,b). Ab1999 uses a kappa LC with 65.8% aa homology with the kappa LC of PCDN.76 UCA (Extended Data Fig. 4b). It has 2 and 14 nt SHMs in the VH and VK genes respectively (Fig. 4a, Extended Data Fig. 4a and Extended Table 2).

**Figure 4:**
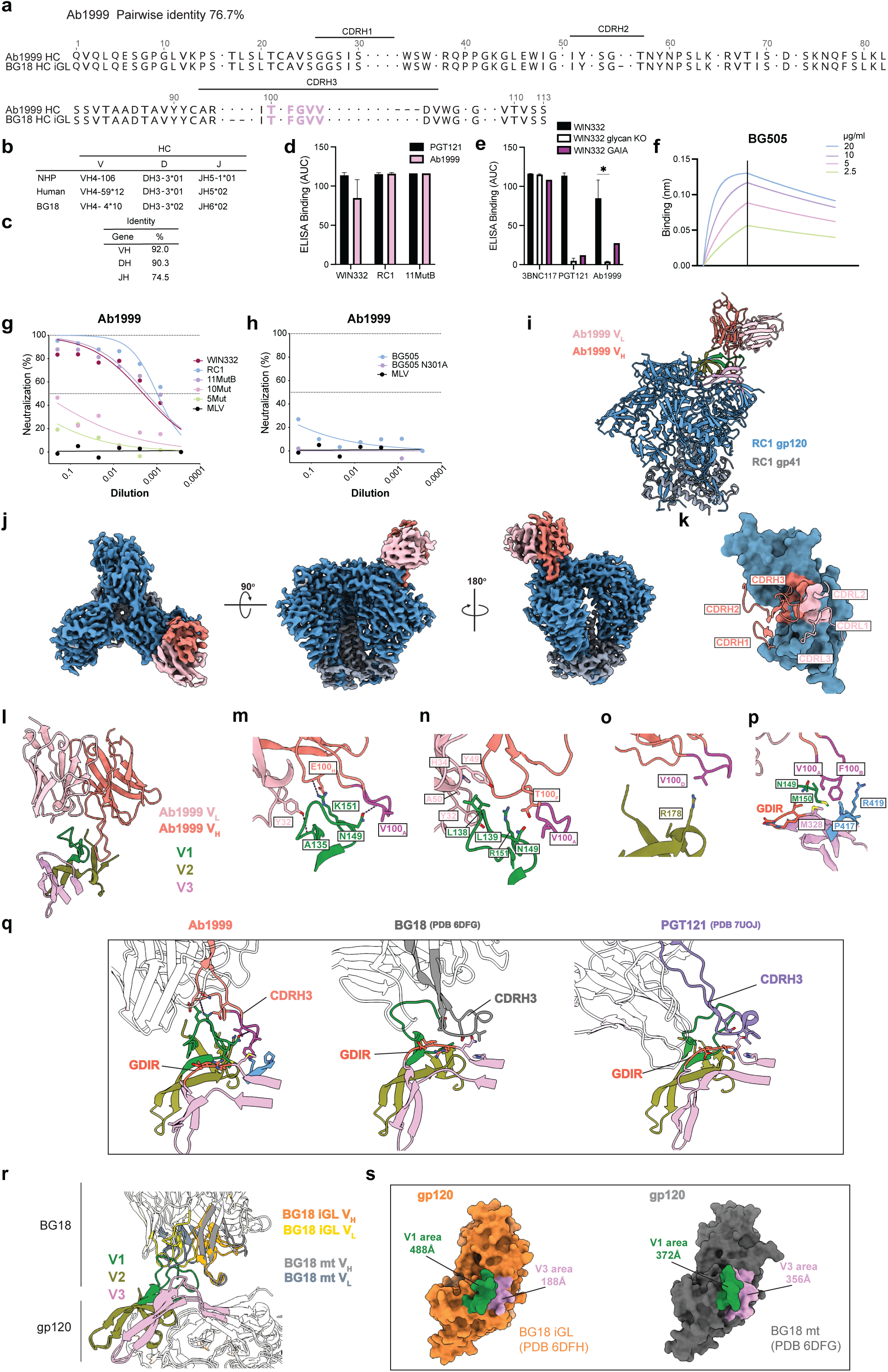
Characterization of Ab1999, a neutralizing BG18-like antibody isolated three weeks after WIN332 prime in NHPs. **a.** Alignment of the amino acid sequences of Ab1999 and inferred germline (iGL) BG18 heavy chains (HC). Dots represent amino acids that are different between sequences. The key TxFGVV CDRH3 motif for neutralization and binding of BG18 is highlighted in pink. Kabat nomenclature is used. **b.** Table shows the VDJ gene assignments for Ab1999 when using macaque and human immunoglobulin gene databases and the human genes of BG18. **c.** Table shows the identity between the VH, DH and JH genes of Ab1999 and BG18. **d.** ELISA binding of Ab1999 to the indicated engineered Envs. **e.** ELISA binding of Ab1999 to WIN332 and two mutant versions of WIN332 that abrogate the V3-glycan epitope: WIN332glycanKO, which differs from WIN332 in that it lacks the N301 glycan; and WIN332 GAIA with a mutated GDIR motif . PGT121 and 3BNC117 were used as controls in d and e at 5µg/ml. **f.** BLI (OCTET) curves showing binding of Ab1999 to fully glycosylated native-like BG505 Env (BG505 T332N). **g.** Results of TZM-bl assays showing neutralization activity of Ab1999 against pseudoviruses carrying Envs with the gradually more native V3-glycan epitopes WIN332, RC1, 11MutB, 10Mut, 5Mut. **h.** Results of TZM-bl assays showing neutralization of Ab1999 against the fully glycosylated BG505 pseudovirus and a mutant version of this that abrogates the V3-glycan epitope (BG505 N301A). Murine Leukemia Virus (MLV) was used as a negative control in g and h. **i.** Atomic model of the RC1 Env-Ab1999 complex, showing interactions between Env (gp120 in blue, gp41 in gray) and Ab1999’s variable domains (salmon and pink). **j.** Cryo-EM density map of the RC1 Env trimer (gp120 in blue and gp41 in gray) bound to a single Ab1999 Fab (Resolution is 3.9 Å). Three views highlight the Fab’s orientation and binding site. **k.** A close-up view highlights the Fab’s CDRs. Gp120 is shown as surface with contact residues colored according to Ab1999 CDR loop. **l.** Overview of the interactions of Ab1999 with the V1 (green), V2 (olive), and V3 (pink) loops. **m-p.** Detailed views of interactions with each loop including hydrogen bonds and hydrophobic interactions. TxFGVV motif is colored in magenta. Antibody residues are numbered using Kabat nomenclature. **m**. Hydrogen bonds (dashed lines) involved in V1 loop contacts with Ab1999 Fab. **n.** Hydrophobic interactions on the V1 loop. **o.** Van der Waal’s contacts on the V2 loop. **p.** Hydrophobic core formed with the tip of TxFGVV motif. **q.** Comparison of the binding interfaces of Ab1999 (salmon), BG18 (gray) and PGT121 (purple) CDRH3s to V1 (green), V2 (olive) and V3 (pink). GDIR motifs are colored in red. **r.** Comparison of BG18 mature (mt) antibody bound to gp120 (gray, PDB 6DFG) with BG18 inferred germline (iGL) precursor antibody bound to gp120 (orange, PDB 6DFH). Zoom-in view of the antibody-gp120 interfaces for both antibodies. **s.** Gp120 surfaces (BG18 mt engaged gp120 in gray and BG18 iGL engaged gp120 in orange) and buried surface areas interacting with BG18 mt and BG18 iGL shown in green for V1 and pink for V3. Measured buried area numbers are labeled acoordingly. Data were analyzed by Student’s t-test and considered statistically significant at *P ≤ 0.05, **P ≤ 0.01, ***P ≤ 0.001 and ****P ≤ 0.0001.

Ab1999 binds to WIN332, RC1 and 11MutB that reintroduce the N332 and the N332 plus N156 glycans respectively (Fig. 4d). Ab1999 does not bind to WIN332glycanKO, a mutant version of WIN332 that abrogates the V3-glycan epitope (Fig. 4e). WIN332glycanKO is different from WIN332 in that it lacks the V3-glycan at position 301, the only signature glycan remaining in WIN332. Ab1999 also shows reduced binding to WIN332 GAIA (Fig. 4e) strongly indicating V3-glycan epitope specificity. Remarkably, Ab1999 binds to fully glycosylated BG505 (Fig. 4f).

To evaluate the capacity of Ab1999 to neutralize HIV, we performed TZM-bl assays against a series of HIV-1 pseudoviruses carrying the autologous WIN332 Env and a series of gradually more native looking Envs including RC1, 11MutB, 10Mut, 5Mut, BG505 and the mutant BG505 N301A to map V3-glycan specificity. We observed high neutralization activity against the WIN332, RC1 and 11MutB pseudoviruses and serially reduced neutralization activity against more native-like pseudoviruses (Fig. 4g). Remarkably, Ab1999 showed low but reproducible neutralization activity against the fully glycosylated BG505 pseudovirus and showed no neutralization for the BG505 N301A pseudovirus and MLV control (Fig. 4h), recapitulating the low activity observed in the serum and consistent with low levels of SHM. As an additional control, a different NHP antibody isolated from a WIN332-primed macaque (Ab2045) did not show any detectable neutralization activity against any of these pseudoviruses (Extended Data Fig.4d).

These data demonstrate that a single immunization with WIN332 elicits antibodies that neutralize HIV-1 pseudoviruses including low levels of neutralization against the fully glycosylated autologous BG505 virus.

### Cryo-EM structure of Ab1999

To elucidate the molecular basis of Ab1999 recognition of Env, we determined a cryo-EM structure of Ab1999 Fab in complex with RC1 at a resolution of 3.9 Å (Fig.4i, j, Extended Data Fig. 5 and Extended Data Table 5). The structure revealed a single Ab1999 Fab bound to RC1, with the interaction predominantly mediated by the antibody’s long CDRH3 loop (Fig. 4j,k). Ab1999 engages the V1, V2, and V3 regions of RC1 (Fig.4l) and we did not observe antibody-glycan interactions (Fig. 4l-p). We observed hydrogen bond (h-bond) interactions between CDRH3 and CDRL1 with the V1 loop. In CDRH3, E100_H_ and V100_A_ interact with K151 and N149, respectively. In CDRL1, Y32 interacts with A135 through a backbone h-bond. (Fig. 4m). We also perceived hydrophobic contacts formed by the CDRL1, CDRL2 and CDRH3 with V1. Y32 and H34 of CDRL1, together with A50 and Y49 of CDRL2 form a hydrophobic patch with L138 and L139 of RC1. In the CDRH3, V100_A_ and T100_F_ interact with N149 and R151 of RC1 (Fig. 4n). Interactions with V2 are less extensive but include Van der Waal’s interactions between V100_D_ of CDRH3 and R178 of RC1 (Fig. 4o). Similarly, Ab1999 engages V3 through the CDRH3 tip via hydrophobic interactions, where residues V100_A_ and F100_B_ of CDRH3 make contacts with M328. In a previous study, the original Q328 residue on V3 was mutated to methionine (Q328M) to increase affinity for BG18 precursors^11^. We observed the CDRH3 tip, especially the TxFGVV motif, engages with a critical hydrophobic core consisting of N149, M150, M328, P417 and R419 (Fig. 4p). Although the CDRH3 of Ab1999 shares the same TxFGVV motif with BG18, the interaction with V3 and the structural comparison with BG18 and PGT121 show a distinct binding mechanism. Notably, Ab1999 binds RC1 by recognizing an alternative hydrophobic pocket consisting of V1, V2, V3 (M328) and C4 without the involvement of the N332 glycan (Fig, 4q).

Since Ab1999 was an early elicited antibody, we compared its epitope with the epitope of BG18 iGL. We compared the Fab-Env structures of iGL and mature BG18 and observed that the buried areas for BG18 iGL at V1 and V3 were 488 Å^2^ and 188 Å^2^, respectively, whereas for mature BG18 they were 372 Å^2^ and 356 Å^2^ (Fig. 4r,s). These data indicated that the BG18 precursor more strongly relied on the V1 loop over the V3 loop for binding and that upon maturation, V1 and V3 dependencies were reduced and increased respectively. At this stage of maturation, the engagement of Ab1999 was similarly more dependent on V1 than V3. In fact, WIN332 and related previously described immunogens include several mutations on V1 introduced to increase affinity for the V3-glycan epitope^11^ (Extended Data Table 1).

We conclude that Ab1999 binds to the V3-glycan epitope of Env in a N332-glycan independent manner.

### WIN332 elicits EPTC112-like antibodies in NHPs

Ab1983 was selected for further characterization for its extraordinarily high similarity with the human type-II V3-glycan bNAb, EPTC112^15^ (Fig. 5a). The HC of Ab1983 has 83.3% aa identity with the HC of iGL EPTC112, 61.9% with the mature EPTC112 HC (Extended Data Fig. 4c) and contains all the aa motifs critical for the neutralization activity of EPTC112^15^ (Fig 5a). These motifs include two serines and one tyrosine in the CDRH1, one serine in the CDRH2 and an “SGW” motif in the CDRH3 (Fig 5a). Ab1983 uses macaque VH4S13, DH6-31, JH4*01, VL9-84 and JL2A*01 genes (Fig 5b). VH4S13 and DH6-31 share 91.7% and 95.2% identity with the human VH4-59 and DH6-19 genes of EPTC112 (Fig.5c). Ab1983 has 3 nt and 1 nt SHMs in the VH and VL genes respectively and a 14-aa-long CDRH3, which is comparable in length to the 15-aa-long CDRH3 of EPTC112 (Fig. 5a and Extended Data Table 2).

**Figure 5:**
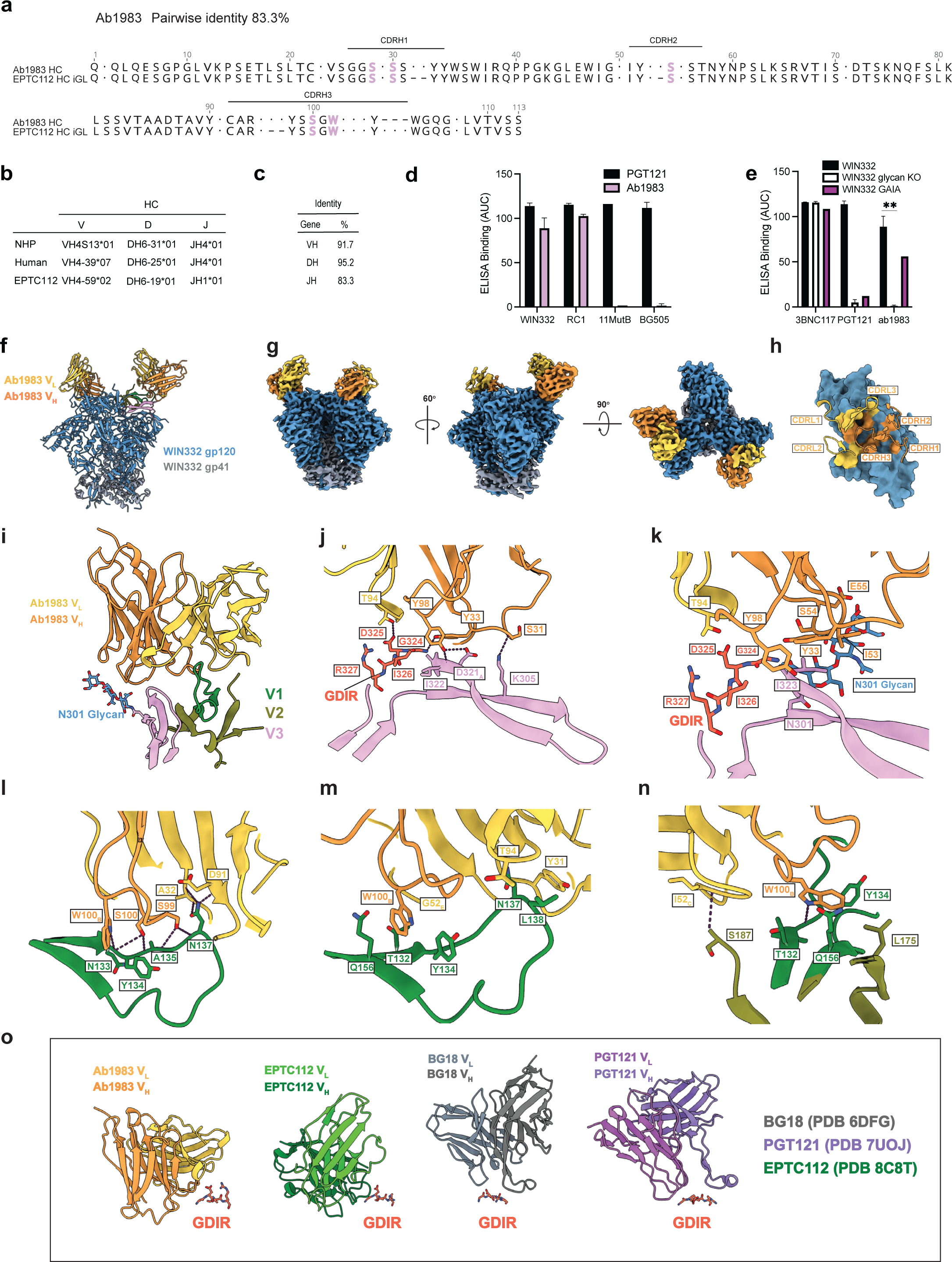
Characterization of Ab1983, an EPTC112-like antibody isolated three weeks after WIN333 prime in NHPs. **a.** Alignment of the amino acid sequences of Ab1983 and inferred germline (iGL) EPTC112 heavy chains (HC). Dots represent amino acids that are different between sequences. Key motifs for neutralization and binding of EPTC112 are highlighted in pink. **b.**Table shows the V_H_D_H_J_H_ gene assignments for Ab1983 when using macaque and human immunoglobulin gene databases and the human genes of EPTC112. **c.** Table shows the homology between the VH, DH and JH genes of Ab1983 and EPTC112. **d.** ELISA binding of Ab1983 to the indicated engineered Envs. PGT121 was used as a positive control at 5µg/ml. **e.** ELISA binding of Ab1983 to WIN332 and two mutant versions of WIN332 that abrogate the V3-glycan epitope: WIN332glycanKO and WIN332 GAIA. **f.** Atomic model of the WIN332 Env-Ab1983 complex. **g.** Cryo-EM density map of the WIN332 Env trimer (gp120 in blue, gp41 in gray) bound to two Ab1983 Fabs (Resolution of 3.8 Å). Three views highlight the Fab’s orientation and binding site. **h.** Close-up view of the Ab1983 paratope, with annotated CDRs that interact with the Env epitope. Gp120 is shown as surface with contact residues colored according to Ab1983 CDR loops. **i.** Interactions of Ab1983 with V1 (green), V2 (olive), and V3 (pink). **j,k.** Detailed hydrogen bonds (j) and hydrophobic interactions (k) with the V3 loop. Hydrogen bonds (dashed lines) between Ab1983 residues and Env residues are shown, along with N301 glycan-mediated interactions. **l,m**. Interactions with V1. Hydrogen bonding (dashed lines) (l) and hydrophobic interactions (m) are shown. **n**. Limited interaction with V2 and the hydrophobic core formed by W100_B_. **o.** Comparison of the mechanisms used by Ab1983, EPTC112, BG18 and PGT121 to bind the GDIR motif. GDIR motifs of are colored in red. Data were analyzed by Student’s t-test and considered statistically significant at *P ≤ 0.05, **P ≤ 0.01, ***P ≤ 0.001 and ****P ≤ 0.0001.

Ab1983 showed high binding to WIN332 and RC1 in ELISA (Fig. 5d) and did not bind to 11MutB, native BG505 and WIN332 glycan KO (WIN332 with a N301-glycan deletion) suggesting that Ab1983 requires the N301 glycan for binding (Fig 5d,e). Ab1983 showed reduced binding to WIN332 GAIA suggesting that antibody binding depends on an intact GDIR motif (Fig.5e).

### Cryo-EM structure of Ab1983

To elucidate the molecular basis of Ab1983 recognition of WIN332, a cryo-EM density map was reconstructed at a resolution of 3.8 Å (Fig 5f-g, Extended Data Fig. 5 and Extended Data Table 5). The structure revealed two copies of Ab1983 bound to WIN332 Env (Fig. 5f-g). Ab1983 Fab engages with V1, V2, and V3 regions, with its heavy chain forming most contacts with V3 (Fig. 5h-j). The Ab1983-WIN332 structure revealed a complex interaction network involving h-bonds and hydrophobic interactions across the V1, V2, and V3 loops. We observed that CDRL3, CDRH1, CDRH2 and CDRH3 loops of Ab1983 interact with V3 through both h-bonds and hydrophobic contacts (Fig5. i, j). In CDRL3, the side chain of T94 forms both hydrogen and hydrophobic interactions with D325. In CDRH1, Y33 engages the G324 of the GDIR motif through an h-bond and I323 through hydrophobic contacts, whereas S31 forms an h-bond with the side chain of K305. In CDRH2, I53 forms hydrophobic contacts with I323. In CDRH3, Y98 forms h-bonds with the backbone of I322 and the side chain of D321_A_, while simultaneously making hydrophobic contacts with I326. (Fig. 5j). Ab1983, similar to EPTC112 and PGT121, binds the N301 glycan, although using different binding mechanisms^15,22^. I53, S54 and E55 in the CDRH2 of Ab1983 are critical for the contact with the N301 glycan (Fig. 5k). Furthermore, the CDRH3, CDRL1, CDRL2 and CDRL3 loops engage with V1 (Fig. 5l,m). Ab1983 forms hydrophobic contacts between W100_B_ of CDRH3 and T132, Y134 and Q156 of V1. Also, hydrophobic interactions are formed between Y31 of CDRL1, G52_E_ of CDRL2 and T94 of CDRL3 and L138, T132 and N137, respectively (Fig. 5l,m). In detail, in CDRH3, W100_B_ binds with the backbone of N133 via an h-bond, as well as interacts with a hydrophobic core consisting of T132, Y134, Q156 and L175 (Fig. 5l-n). In addition, S99 and S100 form h-bonds with the backbone of N133, A135 and N137 of WIN332 (Fig. 5l). In CDRL1 and CDRL3, A32 (CDRL1) and D91 (CDRL3) backbones interact with N137 side chain through h-bond interactions, whereas Y31 (CDRL1) and T94 (CDRL3) form hydrophobic contacts with L138 and N137, respectively. In CDRL2, G52_E_ interacts with T132 via hydrophobic contacts (Fig. 5m). At V2, only limited contacts were formed by the backbone of I52_C_ in the CDRL2 with S187 of WIN332 (Fig. 5n). Notably, Ab1983 engages with the GDIR motif through a combination of h-bonds and hydrophobic interactions between CDRH1 (Y33), CDRH3 (Y98), CDRL3 (T94) and G324, I326 and D325, respectively (Fig. 5j,k). When comparing with other V3-glycan bNAbs, we observed the heavy chain of Ab1983 adopts a similar angle of approach to the GDIR motif as EPTC112, but engages the GDIR motif differently from BG18 and PGT121 (Fig. 5o)

We conclude that Ab1983 is an EPTC112-like antibody that recognizes the V3-glycan epitope in a N332-glycan independent manner and through interactions with V1, V3, and the N301 glycan.

### Sequential immunization boosts the neutralization activity elicited by WIN332 in NHPs

To boost the neutralizing response elicited by WIN332, we sequentially immunized these NHPs with a bolus injection of 7Mut-ST2-N332Y, a modified version of 7Mut^11^ where several glycan holes and the trimer base were masked through the introduction of PNGS sites, and a tyrosine was added at position 332 to contribute a bulky side chain that mimics the presence of the N332 glycan (Fig. 6a). We isolated WIN332-specific GC B cells from dLN collected three weeks after the boost from the sequentially immunized NHPs (Fig. 6b and Extended Data Fig. 6a). Antibody cloning revealed extensive clonal expansion (Fig. 6c) and affinity maturation of WIN332-specific antibodies. Average numbers of mutations were increased to 12.7 nt for HC and 6.9 nt for LC (7.7 nt for VL and 6.2 for VK) (Fig. 6d, e)

**Figure 6:**
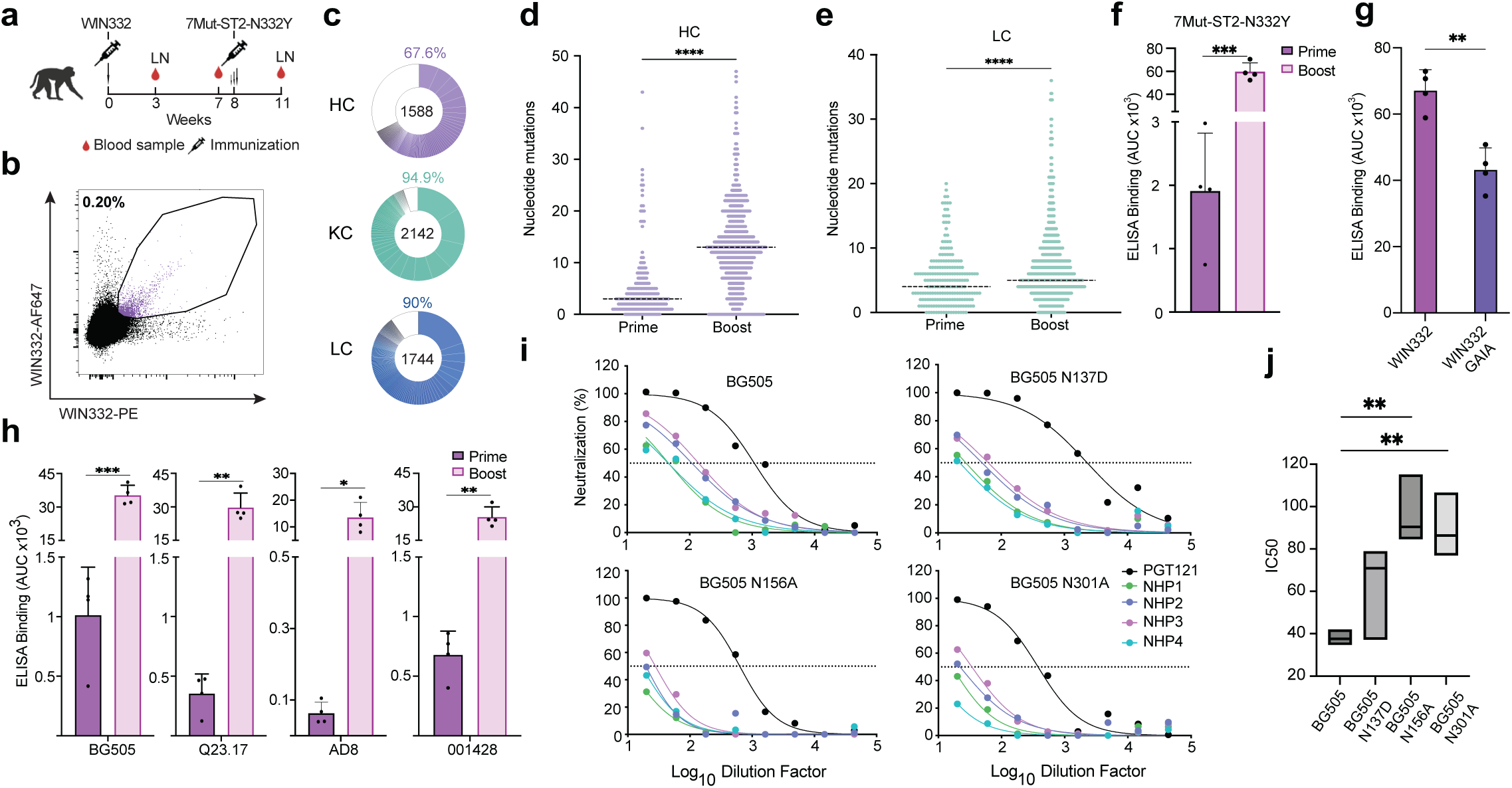
Sequential immunization induces affinity maturation and boosts the autologous neutralization activity in the serum. **a.** Immunization protocol. LN: lymph node biopsy. **b.** Flow cytometry plot showing WIN332-specific B cells (purple) in pooled LN homogenates from sequentially immunized NHPs. **c.** Pie charts showing B cell clonal expansion in sequentially immunized NHPs. Pie charts include heavy chains (HC), kappa light chains (KC) and lambda light chains (LC) from B cells isolated from NHPs 2 and 3. **d, e.** Nucleotide mutations in heavy chains (HC) (d) and light chains (LC) (e) of B cells isolated after WIN332 prime (Prime) and 7Mut-ST2-N332Y boost (Boost). **f.** ELISA binding of the serum collected after WIN332-prime and 7Mut-ST2-N332Y boost to the boosting immunogen 7Mut-ST2-N332Y. **g.** ELISA results showing the binding of the serum collected after the boost to WIN332 and WIN332 GAIA. **h.** ELISA binding of the prime and boost serum to different native Envs. **i.** Results of TZM-bl neutralization assays using purified immunoglobulins from the serum of the sequentially immunized NHPs against the fully glycosylated BG505 pseudovirus (BG505 T332N) and three mutant versions with glycan deletions at the V3-glycan epitope. (BG505 N137D, BG505N156A and BG505 N301A) PGT121 was used as a control. **I.** Graph shows IC50s for the neutralization activity in h. Data were analyzed by Student’s t-test and considered statistically significant at *P ≤ 0.05, **P ≤ 0.01, ***P ≤ 0.001 and ****P ≤ 0.0001.

7Mut-ST2-N332Y strongly boosted the serologic response to the boosting immunogen (Fig. 6f) and the response remained partially dependent on an intact GDIR motif (Fig. 6g). This boost immunization induced significantly stronger binding to the fully glycosylated BG505 trimer and to the heterologous trimers Q23.17, AD8 and 001428 (Fig. 6h)

Remarkably, boost immunization with 7Mut-ST2-N332Y significantly increased the serum neutralization activity against the fully glycosylated BG505 pseudovirus (Fig. 6i) and the neutralization activity was partially specific to the V3-glycan epitope as determined through mapping TZM-bl assays using purified immunoglobulins against the BG505-based mutant pseudoviruses BG505 N137A, BG505 N156A and BG505 N301A and MLV as a control (Fig. 6i, j and Extended Data Fig. 6b)

We conclude that WIN332 followed by sequential immunization with 7Mut-ST2-N332Y elicits neutralizing antibodies that target the V3-glycan epitope of Env.

## Discussion

Sequential immunization is a promising strategy to elicit anti-HIV-1 bNAbs. Important immunogen design efforts are focused on germline-targeting approaches aiming to activate and mature a few selected human bNAb lineages. Several germline-targeting immunogens have been successful at activating precursors of selected human type-I V3-glycan bNAb lineages^1,10,12,14^. Sequential immunization was effective at maturing the precursor of PGT121 into highly mutated, broad and potent PGT121-like antibodies in an immunoglobulin knockin mouse expressing iGL PGT121^1^. Subsequent studies have attempted to achieve neutralization potency and breadth against the V3-glycan epitope through immunization in outbred animals, however, success has been modest, with sporadic elicitation of neutralizing responses of limited potency and breadth^28^ or no neutralization activity^27^.

Reported sequential immunization protocols designed to elicit bNAbs involve numerous immunizations or sustained antigen release over long periods of time^2,13,28,29^. Identifying “premium” antibody lineages that require less convoluted maturation pathways to become bNAbs and designing immunogens that activate them could simplify sequential immunization protocols by reducing the number of immunization steps. Here we used a lineage “agnostic” approach for immunogen design to activate a broad diversity of antibody lineages to the conserved V3-glycan epitope. This approach increases the probability of activating V3-glycan antibody lineages with the potential to mature into bNAbs and broadens the repertoire of antibody lineages to consider for vaccine design purposes.

Human V3-glycan bNAbs show promiscuous binding to the V3-glycan epitope involving different angles of approach and glycan usage, however, all V3-glycan bNAbs isolated until very recently seem to establish key contacts with the N332-glycan^7–9,19^. Multiple observations by us and others led us to hypothesize that there may be antibody lineages that bind the V3-glycan epitope in a N332-glycan independent manner, that N332-glycan dependence may develop during bNAb maturation, and that N332-glycan deficient Env immunogens may be more efficient at activating V3-glycan bNAb precursors. Among our observations, we found antibodies that bound the V3-glycan epitope in a N332-glycan independent manner in previous mouse and NHP studies (unpublished). Moreover, the N332 glycan is underrepresented in transmitted founder viruses (T/F)^30^ and infection with N332-glycan deficient T/F led to the development of N332-glycan dependent V3-glycan bNAbs in two human subjects^30^ suggesting that N332-glycan deficient Env proteins may play a role in bNAb lineage activation. In line with our hypothesis, the UCA of the human V3-glycan bNAb DH270 bound to a synthetic V3-glycopeptide (Man_9_-V3) containing the N332 and N301 glycans but also bound to its aglycone version^31^. Even though DH270 UCA establishes contacts with the N332 glycan in the context of the CH848 d949 Env protein^14^, glycan dependence appears when the antibody is ∼6% mutated^17^ suggesting that the N332-glycan may not be necessary to activate this lineage^32^. Our immunization experiments with WIN332 together with the recent isolation and characterization of a human bNAb that targets the V3-glycan epitope in a N332-glycan independent manner^15^, demonstrate that in addition to type-I N332-glycan dependent lineages, a new class of N332-glycan independent V3-glycan antibodies (type II) should be considered for vaccine design purposes.

WIN332 bound to iGL and UCAs of both type-I and type-II human bNAbs suggesting that WIN332 could be efficient at broadly activating V3-glycan bNAb precursors in humans. In NHPs, WIN332 induced a diverse repertoire of N332-glycan independent V3-glycan antibodies with high homology with human type-I bNAbs like PGT121, BG18 or DH270 and also the recently isolated type-II bNAb EPTC112.

Remarkably, WIN332 elicited serum and monoclonal antibodies that could neutralize the autologous fully glycosylated HIV-1 pseudovirus upon one single immunization and at times as short as 3 weeks after immunization. The neutralization activity observed after the prime is low, not reaching IC5O levels, however, high levels of neutralization activity cannot be expected after one single immunization. This low activity is reproducible under different experimental conditions and recapitulated by a monoclonal antibody (Ab1999) isolated from the macaques. Furthermore, a sequential boost immunization induced affinity maturation of V3-glycan lineages activated by WIN332 and increased the magnitude of the neutralizing activity of the serum. This early binding to the native V3-glycan epitope of Env and neutralization activity has not been previously observed in response to immunization with any reported immunogen or immunization strategy.

Of note, the mAbs isolated from the immunized NHPs, including the neutralizing Ab1999, had very low amounts of SHM, suggesting that moderate levels of SHM could be sufficient to achieve potency and breadth. Ab1999 binding to RC1 Env involves important contacts with V1, which parallels the higher involvement of V1 versus V3 in the mechanism of binding of BG18 iGL and other BG18-like antibodies^12^. The maturation of BG18 resulted in increased contacts of V3 to strengthen the binding to the epitope and reduced engagements of V1 suggesting that a similar maturation path could be possible for Ab1999.

A striking feature of the reported type-II EPTC112 bNAb was its short CDRH3 of 15 aa which is the average length found in the human antibody repertoire. Ab1983, the EPTC112-like NHP antibody characterized here, has a similar CDRH3 of 14 aa. Without the requirement of a long CDRH3, this class of antibody is expected to be easier to elicit through immunization.

Altogether, here we establish two classes of V3-glycan antibody lineages: type-I, N332-glycan dependent and type-II, N332-glycan independent antibodies and report WIN332, the first Env immunogen rationally engineered to activate and mature antibody lineages that do not require the N332-glycan at early stages of maturation. The early neutralization induced by WIN332, presents this immunogen as a promising candidate to shorten sequential immunization protcols and streamline HIV-1 vaccine development.

## Methods

### Envelope proteins

The newly-engineered Env SOSIP trimers, WIN332, WIN332-GAIA-Avitag, WIN332-Avitag, 7MUT-ST2-N332Y and other previously reported engineered SOSIP Env trimers, RC1, RC1-glycan knockout (same as WIN332 glycan KO)^10^, 11MUTB, 10MUT, 7MUT, 5MUT^21^ and native-like BG505 T332N^16^, CEO217, Q23, AD8, BJOX2000, CNE55, 001428, CH119, AMC011, were cloned in the pPPI4 or pVAX expression vectors using synthetic gene fragments (Integrated DNA technologies (IDT)) or were produced by Genescript. Specific mutations for each protein are listed in Extended Data Table 1.

Env trimers were expressed as soluble native-like gp140 trimers^16^. Soluble Env trimers were expressed by transient transfection in Expi293 cells (Thermo Fisher Scientific) and purified from cell supernatants by Lectin chromatography and size exclusion chromatography (SEC) as previously described^33^. Proteins were stored at -80°C in PBS until the time of use.

### Analysis of Site-Specific N-linked Glycopeptides by Liquid chromatography-Mass Spectrometry (LC-MS)

Aliquots of purified WIN332 protein were reduced by incubating with 10 mM dithiothreitol (Sigma) at 56 °C and alkylated by 27.5 mM iodoacetamide (Sigma) at room temperature (RT) in dark. The aliquots were then digested respectively by using a combination of alpha lytic protease (New England BioLabs), AspN (Promega), chymotrypsin (Athens Research and Technology), Lys-C (Promega), Glu-C (Promega), and trypsin (Promega). The resulting peptides were separated on an Acclaim™ PepMap™ 100 C18 column (75 μm x 15 cm) and eluted into the nano-electrospray ion source of an Orbitrap Eclipse™ Tribrid™ mass spectrometer (Thermo Scientific) at a flow rate of 200 nL/min. The elution gradient consists of 1-40% acetonitrile in 0.1% formic acid over 370 min followed by 10 min of 80% acetonitrile in 0.1% formic acid. The spray voltage was set to 2.2 kV and the temperature of the heated capillary was set to 275 °C. Full MS scans were acquired from m/z 200 to 2000 at 60k resolution, and MS/MS scans following higher-energy collisional dissociation (HCD) with stepped collision energy (15%, 25% , 35%) were collected in the orbitrap at 15k resolution. pGlyco3^34^ was used for database searches with mass tolerance set as 20 ppm for both precursors and fragments. The database search output was filtered to reach a 1% false discovery rate for glycans and 10% for peptides. The glycan and peptide assignment for each spectra was then manually validated after filtering. Quantitation was performed by calculating spectral counts for each glycan composition at each site. Any N-linked glycan compositions identified by only one spectra were removed from quantitation. N-linked glycan compositions were categorized into 19 classes (including “Unoccupied” as class 19): HexNAc(2)Hex(9∼5)Fuc(0∼1) was classified as M9 to M5 respectively; HexNAc(2)Hex(4∼1)Fuc(0∼1) was classified as M1-M4; HexNAc(3∼6)Hex(5∼9)Fuc(0)NeuAc(0∼1) was classified as Hybrid with HexNAc(3∼6)Hex(5∼9)Fuc(1∼2)NeuAc(0∼1) classified as F-Hybrid; Complex-type glycans are classified based on the number of antenna and fucosylation: HexNAc(3)Hex(3∼4)Fuc(0)NeuAc(0∼1) is assigned as A1 with HexNAc(3)Hex(3∼4)Fuc(1∼2)NeuAc(0∼1) assigned as F-A1; HexNAc(4)Hex(3∼5)Fuc(0)NeuAc(0∼2) is assigned as A2/A1B with HexNAc(4)Hex(3∼5)Fuc(1∼5)NeuAc(0∼2) assigned as F-A2/A1B; HexNAc(5)Hex(3∼6)Fuc(0)NeuAc(0∼3) is assigned as A3/A2B with HexNAc(5)Hex(3∼6)Fuc(1∼3)NeuAc(0∼3) assigned as F-A3/A2B; HexNAc(6)Hex(3∼7)Fuc(0)NeuAc(0∼4) is assigned as A4/A3B with HexNAc(6)Hex(3∼7)Fuc(1∼3)NeuAc(0∼4) assigned as F-A4/A3B; HexNAc(7)Hex(3∼8)Fuc(0)NeuAc(0∼1) is assigned as A5/A4B with HexNAc(7)Hex(3∼8)Fuc(1∼3)NeuAc(0∼1) assigned as F-A5/A4B.

### Analysis of Deglycosylated HIV-1 envelope protein by LC-MS

Aliquots of the purified WIN332 protein were reduced by incubating with 10 mM dithiothreitol (Sigma) at 56 °C and alkylated by 27.5 mM iodoacetamide (Sigma) at RT in dark. The aliquots were then digested respectively using chymotrypsin (Athens Research and Technology), Glu-C (Promega), Lys-C (Promega), Arg-C (Promega), and trypsin (Promega). Following digestion, the extracted peptides were deglycosylated by Endoglycosidase H (Promega) followed by PNGaseF (Promega) treatment in the presence of 18O water (Cambridge Isotope Laboratories). The resulting peptides were separated on an Acclaim™ PepMap™ 100 C18 column (75 μm x 15 cm) and eluted into the nano-electrospray ion source of an Orbitrap Eclipse™ Tribrid™ mass spectrometer (Thermo Scientific) at a flow rate of 200 nL/min. The elution gradient consists of 1-40% acetonitrile in 0.1% formic acid over 370 min followed by 10 min of 80% acetonitrile in 0.1% formic acid. The spray voltage was set to 2.2 kV and the temperature of the heated capillary was set to 275 °C. Full MS scans were acquired from m/z 200 to 2000 at 60k resolution, and MS/MS scans following collision-induced dissociation (CID) at 38% collision energy and higher-energy collisional dissociation (HCD) with stepped collision energy (15%, 25%, 35%) were collected in the ion trap. The spectra were analyzed using SEQUEST (Proteome Discoverer 2.5, Thermo Fisher Scientific) as well as Byonic (v4.1.10, Protein Metrics) ^35^ with mass tolerance set as 20 ppm for precursors and 0.5 Da for fragments. The search output was filtered to reach a 1% false discovery rate at the protein level and 10% at the peptide level. The site assignment for each spectra was then manually validated after filtering. Occupancy of each N-linked glycosylation site was calculated using spectral counts assigned to the 18O-Asp-containing (PNGaseF-cleaved) and/or HexNAc-modified (EndoH-cleaved) peptides and their unmodified counterparts.

### Animals

Rhesus macaques (*Macaca mulatta*) of Indian genetic origin, 4 years of age, were housed and cared for in a biosafety level 2 facility at the Tulane National Primate Research Center (TNPRC). All animal procedures and experiments were performed according to protocols approved by the TNPRC IACUC. SMNP was produced as previously described^20^. At day 0, macaques received one subcutaneous bolus immunization with a total of ∼200 μg of soluble WIN332 SOSIP trimer adjuvanted in 375 μg of SMNP distributed between the 4 legs ( ∼50 µg per foreleg and ∼50µg per hind leg). Macaques were boosted with a total amount of 200 μg of soluble 7MUT-ST2-N332Y in 375 μg of SMNP adjuvant using an escalating dose delivery approach^36^ at week 8 (1 dose of 30 µg of WIN332 in 56.5 µg SMNP at day 1, 1 dose of 60 µg of WIN332 in 112 µg SMNP at day 3 and 1 dose of 110 µg of WIN332 in 206.5 µg SMNP at day 5) and distributed between the 4 legs like the prime. Blood samples were obtained from the macaques prior to immunization, 3 and 7 weeks after WIN332 prime and 3 weeks after 7MUT-ST2-N332Y boost immunization (week 11). Lymph node biopsies were obtained from the immunized macaques 3 weeks after WIN332 prime and 3 weeks after boost immunization with 7MUT-ST2-N332Y.

### ELISA

ELISAs with SOSIP Env trimers WIN332, RC1, WIN332glycan KO, 11MUTB, 10MUT, 7MUT, 5MUT and native Env trimers were performed as previously described^10^. Briefly, 96-well plates were coated with 50 μl of a solution of the trimer at 4 μg/ml in 1xPBS (Corning # 20-030-CV) and incubated at 4°C overnight. Plates were washed with washing buffer (1x PBS with 0.05% Tween-20 (Sigma-Aldrich # P7949)) and blocked (1x PBS with 5% milk) for 1 hour (h) at RT. Serum samples were assayed at a 1:100 or 1:30 starting dilution, supernatants from Expi293 cells transfections were assayed at a 1:3 starting dilution and the antibody controls were added at 5μg/ml all followed by seven additional 3-fold serial dilutions for 1h at RT. Plates were washed three times and incubated with an anti-human IgG secondary antibody conjugated to Horse Radish Peroxidase (HRP) (Jackson ImmunoResearch #115-035-071) at 1:5000 dilution in washing buffer for 1 h at RT. Plates were developed by adding ABTS and Optical Density (OD) was measured at 405 nm.

Alternatively, 96-well plates were coated with 50 μl of a solution of PGT121 or 3BNC117 antibodies engineered to have a mouse Fc at 4 μg/ml in 1xPBS and incubated at 4°C overnight. Plates were washed with washing buffer (1x PBS with 0.05% Tween-20), blocked (1x PBS with 5% milk) for 1 h at RT and incubated in 50 μl of a solution of an Env trimer at 4 μg/ml in blocking buffer for 1 h at RT. Plates were washed three times and incubated with 5 μg/ml of an antibody with human Fc and 3-fold serial dilutions for 1 h. Plates were washed three times and incubated with an anti-human IgG secondary antibody conjugated to Horse Radish Peroxidase (HRP) at 1:5000 dilution in washing buffer. Plates were developed by adding ABTS and Optical Density (OD) was measured at 405 nm.

For competition ELISAs, 96-well plates were directly coated with 50 μl of a solution of an Env trimer at 4 μg/ml in 1xPBS and incubated at 4°C overnight. Plates were washed three times in washing buffer and blocked in blocking buffer for 1 h at RT. Plates were then incubated in 50 μl of a 15 μg/ml solution of PGT121 or 3BNC117 with a mouse Fc in blocking buffer for 1.5 h at RT. Plates were washed three times and NHP serum or mAb were added at the indicated starting dilutions and six additional 3-fold serial dilutions for 30min at RT. Plates were washed and developed using anti-human IgG secondary antibody conjugated to HRP at 1:5000 dilution in washing buffer. Plates were developed by adding ABTS and Optical Density (OD) was measured at 405 nm.

Endpoint titers were calculated as the reciprocal number of the dilution at which the sample OD was 3 times the background.

### Flow cytometry and single B-cell sorting

Frozen cell homogenates from lymph node (LN) biopsies obtained from immunized macaques were thawed and washed in RPMI medium 1640 (1x) (Gibco # 11875-093).

LN cells were incubated with 100 μl of FACS buffer (PBS 1x with 2% fetal bovine serum and 1mM EDTA) with human Fc Block (BD Biosciences # 564219) at a 1:500 dilution for 30 min on ice.

WIN332, WIN332-GAIA and BG505 tetramers were prepared by incubating 5 μg of Avitagged and biotinylated WIN332 (WIN332-AviBio), WIN332-GAIA (WIN332-GAIA-AviBio) or BG505 (BG505-AviBio) with fluorophore-conjugated streptavidin at a 1:200 dilution in 1xPBS for 30 min on ice. WIN332^++^ WIN332-GAIA^-^ or BG505**^++^** LN B-cells were isolated using WIN332-AviBio or BG505-AviBio conjugated to streptavidin AF647 (BioLegend, # 405237), WIN332-AviBio or B41-AviBio conjugated to streptavidin PE (BioLegend, # 405204) and WIN332-GAIA-AviBio conjugated to streptavidin BV605 (BioLegend, # 405229) as baits. Tetramers were mixed with the antibody cocktails indicated below to a final concentration of 5 μg/ml each.

Macaque LN cells were stained with: anti-CD3 APC-eFluor 780 (Invitrogen # 47-0037-41), anti-CD14 APC-eFluor 780 (Invitrogen # 47-0149-42), anti-CD16-APC-eFluor 780 (Invitrogen # 47-0168-41), anti-CD8 APC-eFluor 780 (Invitrogen # 47-0086-42), anti-CD20 PE-Cy7 (BD biosciences # 335793), and anti-CD38 FITC (Stem Cell # 60131FI) at 1:200 dilution and the LIVE/DEAD marker Zombie NIR (Biolegend # 423106) at 1:400 dilution. Cells were incubated with the mixture of tetramers and antibody cocktail for 30 min on ice.

Zombie NIR^-^/CD16^-^/CD8a^-^/CD3^-^/CD14^-^/CD20^+^/CD38^+^/WIN332^++^/WIN332-GAIA^-^ or BG505^++^/WIN332-GAIA^-^ single cells were isolated from the macaque LN cell homogenates using a FACS Aria III (Becton Dickinson). Single B cells were sorted into individual wells of 96-well plates containing 5 μl of lysis buffer (TCL buffer (Qiagen #1031576) with 1% of 2-β-mercaptoethanol). The cell lysates were stored at −80°C or immediately used for subsequent mRNA purification. Alternatively, trimer-specific B cells were sorted in RPMI with 5% FBS for 10X genomics.

### Antibody sequencing

RNA was purified from single cells using paramagnetic beads (2.2x ratio) (RNAClean XP, # A63987 Beckman Coulter). RNA was eluted from the magnetic beads with 11 μl of a solution containing (14.5 ng/μl of random primers (Invitrogen, # 48190-011), 0.5% of tergitol, (Type NP-40, 70% in H_2_O, Sigma-Aldrich, # NP40S-100ML), 0.6 U/μl of RNAse inhibitor (Promega # N2615) in nuclease free water (Qiagen), and incubated at 65°C for 3 min. cDNA was synthesized by reverse transcription (SuperScript® III Reverse Transcriptase, Invitrogen, # 18080-044, 10’000U)^37^. cDNA was stored at - 20°C or 10 μl of nuclease free water were added to the cDNA before subsequent use for antibody gene amplification by nested Polymerase Chain Reaction (PCR).

Macaque IgH and IgK/L genes were amplified and cloned^37^ using the primers and PCR protocols in Extended Data Tables 3 and 4^38^. Briefly, after a first PCR reaction to amplify the antibody genes, a second PCR (cloning PCR) introduced nucleotide tails that were used for subsequent Ligation Independent Cloning (LIC) in expression vectors^38^ (see antibody cloning). The cloning PCR products were sequenced by Sanger sequencing using the primers in Extended Data Table 3.

### 10X Genomics

Single WIN332-specific GC B cells were bulk-sorted in RPMI with 5% FBS, processed by Wistar Genomics Core following the manufacturer protocol (Chromium Next GEM Single Cell V(D)J protocol v3) with modifications^39^and macaque V(D)J libraries were generated.

Illumina NGS-generated raw data files were processed using the Cell Ranger single cell gene expression software provided by 10X Genomics. Sequences were quality filtered, followed by assembly and annotation using Cell Ranger adapted to contain rhesus macaque Ig genes from a custom reference. Contigs assembled by Cell Ranger were reannotated. To assign antibody sequences to each barcode, a custom Bash pipeline was developed to generate a CSV file that organizes query sequences by heavy chain (VH), light chain kappa (VK), and light chain lambda (VL). This pipeline incorporates a Python script that automatically aligns query sequences to germline V(D)J segments using the NCBI IgBLAST database. The resulting annotations are parsed and structured into separate columns based on their corresponding variable regions (VH, VK, VL). Finally, sequences sharing the same identifier are matched and paired by row to represent complete antibody repertoires. After pairing, sequences were analyzed with a custom pipeline with macaque Ig libraries to determine B cell immunogenetics and clonality ^40^.

### Antibody cloning

Antibody cloning was performed by Genescript or in house through LIC cloning. For LIC cloning, the cloning PCR products were purified using the PCR clean-up kit (Qiagen, 28183). The purified products were cloned into the corresponding expression vectors encoding human IgG, IgL and IgK constant regions digested with the restriction enzymes AgeI and SalI for IgH, AgeI and XhoI for IgL and AgeI and BsiwI for IgK as previously described^38^

### Antibody production and purification

Macaque IgGs were produced by Genescript or in house through transient transfection in Expi293T cells. For transfections, DNA mixtures containing IgH and IgL/IgK antibody genes at 1:1 ratio were used for transfections following manufacturer instructions (Thermo Fisher Scientific, # A14635). IgGs were purified from the supernatants of transfected Expi293T cells using Protein G (Cytiva, cat # 28-4083-47).

For structural studies, macaque Fab was obtained by digesting IgG at 1-5 mg/ml with ficin (Sigma). Fab was purified by SEC chromatography^41^, followed by monoQ 5/50 (GE Healthcare) ion exchange chromatography.

Serum Igs were purified from 200 μl of macaque serum using Ab Spin Trap Protein G Sepharose columns (GE Healthcare, #28-4083-47). Ig-containing fractions were buffer exchanged with PBS using a 30 kDa molecular weight cutoff concentrator (Milipore Sigma, cat #UFC903024).

### *In vitro* neutralization assay

Env-pseudotyped viruses were produced following the standardized protocol ^42^. Briefly, HEK 293T cells were seeded in 100 mm plates at a density of 4 × 10⁶ cells per plate and incubated overnight in complete growth medium. Cells were transfected using FuGENE 6 (Promega) with 4 µg of Env plasmid and 8 µg of backbone plasmid per plate. After a brief incubation (4–5 hours), the medium was replaced, and the culture was maintained for 48 hours. Virus-containing supernatants were harvested, supplemented with fetal bovine serum to a final concentration of 20%, filtered through 0.45 µm membranes, aliquoted into cryovials, and stored at −80°C.

Pseudovirus titers were determined using a tissue culture infectious dose (TCID) assay in TZM-bl cells, a reporter cell line engineered to express luciferase under control of the HIV-1 Tat protein ^42^. Serial 5-fold dilutions of the pseudovirus stock were prepared in 96-well plates, and 10,000 TZM-bl cells per well were added in the presence of DEAE-dextran (Final concentration of 40 µg/mL). After 48 hours of incubation at 37°C, luminescence was measured using the Firefly Luciferase Assay Kit (Merck). Virus input was considered optimal when luminescence signals fell within the range of 15,000 to 50,000 relative luminescence units (RLU).

TZM-bl assays were performed as described^43^. For the assay setup, 96-well flat-bottom plates were used. To reduce edge effects and minimize variability due to evaporation, all peripheral wells (Columns 1, 11, and 12; rows A and H) were filled with sterile phosphate-buffered saline (PBS) and were not used for experimental conditions. The assay was conducted using the remaining internal wells. Column 2 served as the virus control (Cells and virus without antibody), while columns 3 through 10 were used for serial dilutions of the test antibody or serum.

Serial 3-fold dilutions of the test antibody or purified IgGs from macaque serum were performed in duplicate, using wells from bottom to top (Row G to row B) with a starting concentration of 100 µg/mL for WIN332, RC1, 11MutB pseudoviruses, 200 µg/mL for 10Mut and 7Mut and 400 µg/mL for 5Mut, BG505 T332N, BG505 N301A and MLV. Serial 3-fold dilutions of purified IgGs from macaque serum were performed in duplicate, using wells from bottom to top (Row G to row B) with a starting concentration of 300 µg/mL. After dilutions were prepared, 50 μL of pseudovirus diluted in growth medium was added to all experimental wells (columns 2–10).

Plates were incubated at 37°C for 1 h to allow virus-antibody interaction. During this time, a suspension of TZM-bl cells was prepared at 1×10⁵ cells/mL in growth medium containing DEAE-dextran (Final concentration of 40 µg/mL). 100 μL of this suspension was then added to each well. Plates were incubated for 48 h at 37°C in a humidified 5% CO₂ atmosphere.

Neutralization activity was calculated as a function of the reduction in Tat-induced luciferase expression.

### Biolayer interferometry (BLI, OCTET) binding studies

BLI experiments were performed using the OCTET Red96 system to determine affinities of antibodies (Fabs) for WIN332. Biotinylated WIN332–Avitag (WIN332-AviBio) was immobilized on high-precision streptavidin (SAX) biosensors (FORTÉBIO) using a solution of WIN332-AviBio at 5 µg/mL in dilution buffer (FORTÉBIO). Four serial dilutions (20, 10, 5 and 2.5 µg/mL) of each experimental Fab or antibody and PGT121 antibody were prepared in dilution buffer (FORTÉBIO). Alternatively, the experimental antibody (5 µg/mL) was immobilized on Anti-Human Fc Capture (AHC) Biosensors (FORTÉBIO) and assayed against four serial dilutions (20, 10, 5 and 2.5 µg/mL) of WIN332. The binding experiment was performed at 30 °C using the following protocol: baseline 1 (60 s), WIN332 or antibody/Fab loading (300 s), baseline 2 (200 s), association (300 s) and dissociation (600 s). Analysis was performed using OCTET software Data Analysis HT 10.0 (FORTÉBIO).

### EMPEM

Serum from each immunized macaque was 10 fold diluted in PBS. Diluted sera were incubated with protein G sepharose 4 Fast Flow resin (Cytiva) overnight at 4°C. Polyclonal IgG was purified by eluting in 0.1M Glycine-HCl (pH 3.5) and neutralized immediately in 1M Tris-HCl (pH 8). Purifed polyclonal IgG samples were further digested with 2% papain (Thermofisher). 2 mg of digested polyclonal Fab was incubated with 15 µg of WIN332 or RC1 trimer on ice or overnight at 4°C. The complexes were purified by size exclusion chromatography (SEC) through Superose 6 Increase 10/300 GL column (Cytiva). Fractions corresponding to trimer/Fab complexes were concentrated. 4

µL of purified complexes at 0.01 mg/mL were applied to glow-discharged, carbon-coated 400 mesh cooper grid (Electron Microscopy Sciences, EMS). After blotting extra sample, grids were stained with 2% uranyl formate for 2 min followed by blotting extra staining buffer. Grids were imaged in 100kV FEI Tecnai T12 scope at 57,000 magnification equipped with Gatan CCD camera. Collected micrographs were processed in Relion^44^. Specifically, LoG picked particles were cleaned up by 2D classification. Subset classes showing clear trimer or Fab bound trimer were selected for 3D classification. Data analysis and figure generation were performed in UCSF Chimera^45^.

### Cryo-EM sample and grid preparation

RC1 and Ab1999 Fab or WIN332 and Ab1983 Fab were mixed together using a molar ratio of 1:9 (Env:Fab) and incubated for 1 h on ice. Complexes were purified by SEC through Superose 6 increase 10/300 GL column (Cytiva). Fractions containing the Env/Fab complexes were pooled and concentrated down using 100 kDa molecular weight cutoff concentrators. Samples were flash frozen in liquid nitrogen and stored in -80°C.

Complexes were thawed and diluted to around 0.085 g/L. UltrAuFoil 1.2/1.3 300 mesh grids were glow discharged for 1 min and immediately used for graphene oxide application using previously published protocol^46^. Complexes were vitrified by applying 4 μL of the diluted sample to the GO-coated grids using vitrobot Mark IV (FEI, Waltham, MA) at 4°C and 100% humidity followed by plunging into liquid ethane.

### Cryo-EM data collection and processing

Cryo-EM grids containing RC1/Ab1999 or WIN332/Ab1983 complexes were loaded on to a 200 kV Thermo Fisher Glacios Microscope equipped with a Falcon 4 detector. Dose-fractionated movies were collected using a total dose 40 e-/Å2 across 40 frames or 50 e-/Å2 across 49 frames for WIN332/Ab1983 or RC1/Ab1999, respectively. Both datasets were collected with a defocus range of -0.8 to -2.2 μm.

Data processing for both datasets was done similarly. In summary, raw movies were imported into CryoSPARC and preprocessed using Patch Motion Correction and Patch CTF Estimation. Particles were picked using blob picker and then extracted with binning by 4 resulting in pixel size of 3.79 Å/pixel. Particles were first cleaned up using 2D Classification and particles in the junk classes were discarded. Remaining particles were used for Ab initio reconstruction and the resulting 3D volumes were used for subsequent Heterogeneous Refinement. Particles belonging to the good 3D classes were selected and reextracted with binning by 2. This was followed by another round of Heterogeneous Refinement. The selected particles were then reextracted with the original pixel size and used for Non-Uniform Refinement. The resulting auto-sharpened maps were used for model building and analysis.

### Model Building

Initial models of Ab1999 and Ab1983 Fabs were generated using SWISS-MODEL^47^ without reference structures. The previously published model of RC1 (PDB: 6ORO) was used as the initial model for model building for both RC1/Ab1999 and WIN332/Ab1983. WIN332 Env model was modified by deleting the N332 glycan and mutating the asparagine intoglutamine. Reference models were docked into the density maps in UCSF ChimeraX^48^ and manually real-space refined in Coot^49^. N-linked glycans were added to the models manually in Coot^49^ . FModels were refined in Rosetta and real-space refined in Coot iteratively. Model geometry was validated by MolProbity^50^, glycan conformation was validated by Privateer^51^, and model fit-to-map was validated by EMRinger^52^.

### Analysis

Geneious Prime was used for sequence analysis. Flow cytometry data were processed using FlowJo v.10.5.0. GraphPad Prism 10 was used for data analysis. Structural analysis was performed using the sowtware indicated in the sections above. For antibody analysis, a custom V(D)J database was used with modifications^40^.

### Quantification and statistical analysis

Statistical information, including sample size (n), mean, standard deviation and statistical significance values are indicated in the text or the figure legends. GraphPad Prism 10 was used for statistical analysis by paired Student’s t-test. Data were considered statistically significant at *P ≤ 0.05, **P ≤ 0.01, ***P ≤ 0.001 and ****P ≤ 0.0001.

### Data availability

Nucleotide sequences of relevant macaque antibodies will be deposited at GenBank after acceptance of the manuscript. Cryo-EM maps and models have been deposited with accession codes: (EMD-70231 and EMD-70395). The mass spectrometry data have been deposited to the ProteomeXchange Consortium via the MassIVE partner repository (https://massive.ucsd.edu/ProteoSAFe/static/massive.jsp) with the dataset identifier MSV000097767.

### Code availability

Computer code for macaque immunoglobulin gene analysis is publicly available on GitHub https://github.com/stratust/igpipeline/tree/igpipeline2_timepoint_v2 and https://github.com/MartaTarquis/blastN_pairs.

## Acknowledgments

We thank members of the Escolano, Pallesen, Kulp and Weiner laboratories for valuable discussions, Jeffrey Faust, John Fundyga and Sonali Majumdar at The Wistar’s Flow Cytometry and Genomics cores for assistance. Dr. Hugo Mouquet for providing the nucleotide sequence of EPTC112 and advice to infer its germline precursor. We thank all the authors for critically reading the manuscript and providing valuable feedback.

This work was supported by the National Institute of Allergy and Infectious Diseases (NIAID) of the National Institutes of Health (NIH) Grant R00 AI140770-03 (A.E), 5 P30 AI045008-23 (A.E.), P30 AI045008-24 (A.E), SAP# 4100090384 (A.E), R01 AI172627-01A1 (A.E) and by the Gates Foundation INV-036995 (A.E). J.P. is supported by 5 U19 AI166916-03. D.B.W. is supported by U19 AI166916 , BEAT-HIV UM1AI64570, and the a W.W. Smith Charitable Trust Distinguished Professorship in Cancer Research. Mass spectrometry studies were supported by the National Institutes of Health grant R01GM130915 (L.W.) and R01AI157854 (L.W.), and National Science Foundation grant NSF Biofoundry: Glycoscience Research, Education, and Training Grant 2400220 (L.W.).

Additional support included the Ching Jer Chern Postdoctoral Fellowship (I.R.R).

## Author contributions

I.R.R designed, performed and analyzed experiments, and coordinated all experimental aspects of the project. J.D., Z.L., J.C. and J.P. designed, performed and analyzed Cryo-EM and EMPEM experiments. M.K., M.T-M., E.U., C.B., S.G. designed and performed experiments, M.W. immunized rhesus macaques, collected and prepared tissue and cell samples, P.Z. designed, performed and analyzed mass spectrometry experiments, R.H. provided proteins and advice. A.A.W. and M.B.M produced SMNP adjuvant. G.M.S and B.H.H provided UCA and iGL antibodies, pseudoviruses, a macaque immunoglobulin gene reference and valuable advice for TZM-bl neutralization assays. D.I provided SMNP adjuvant, L.W. supervised mass spectrometry experiments, D.W. supervised experiments, reviewed specific serological analysis with S.G. and provided valuable reagents, R.S.V. supervised macaque treatments and sample collection, D.W.K. supervised the design and production of 7MUT-ST2-N332Y, provided valuable reagents and advice, J.P. supervised Cryo-EM and EMPEM studies, A.E. conceived the study, designed WIN332, supervised and coordinated the study. Writing original draft: I.R.R. and A.E. with contributions from P.Z., L.W. J.D., Z.L. and J.P. Draft reviewing and editing: all authors.

## Materials & Correspondence

Questions and requests for reagents should be directed to Amelia Escolano.

## Inclusion & Ethics

Research has been conducted following the recommendations set out in the Global Code of Conduct for Research in Resource-PoorSettings including local researchers throughout the research process.

## References

1. Escolano, A., et al. Sequential Immunization Elicits Broadly Neutralizing Anti-HIV-1 Antibodies in Ig Knockin Mice. Cell 166, 1445–1458 e1412 (2016).

2. Kong, R., et al. Antibody Lineages with Vaccine-Induced Antigen-Binding Hotspots Develop Broad HIV Neutralization. Cell 178, 567–584 e519 (2019).

3. Caniels, T.G., et al. Germline-targeting HIV vaccination induces neutralizing antibodies to the CD4 binding site. Sci Immunol 9, eadk9550 (2024).

4. Landais, E., et al. Broadly Neutralizing Antibody Responses in a Large Longitudinal Sub-Saharan HIV Primary Infection Cohort. PLoS Pathog 12, e1005369 (2016).

5. Rusert, P., et al. Determinants of HIV-1 broadly neutralizing antibody induction. Nat Med 22, 1260–1267 (2016).

6. Kong, L., et al. Supersite of immune vulnerability on the glycosylated face of HIV-1 envelope glycoprotein gp120. Nat Struct Mol Biol 20, 796–803 (2013).

7. Freund, N.T., et al. Coexistence of potent HIV-1 broadly neutralizing antibodies and antibody-sensitive viruses in a viremic controller. Sci Transl Med 9 (2017).

8. Mouquet, H., et al. Complex-type N-glycan recognition by potent broadly neutralizing HIV antibodies. Proc Natl Acad Sci U S A 109, E3268–3277 (2012).

9. Garces, F., et al. Structural evolution of glycan recognition by a family of potent HIV antibodies. Cell 159, 69–79 (2014).

10. Escolano, A., et al. Immunization expands B cells specific to HIV-1 V3 glycan in mice and macaques. Nature 570, 468–473 (2019).

11. Steichen, J.M., et al. HIV Vaccine Design to Target Germline Precursors of Glycan-Dependent Broadly Neutralizing Antibodies. Immunity 45, 483–496 (2016).

12. Steichen, J.M., et al. A generalized HIV vaccine design strategy for priming of broadly neutralizing antibody responses. Science 366(2019).

13. Saunders, K.O., et al. Vaccine Elicitation of High Mannose-Dependent Neutralizing Antibodies against the V3-Glycan Broadly Neutralizing Epitope in Nonhuman Primates. Cell Rep 18, 2175–2188 (2017).

14. Saunders, K.O., et al. Targeted selection of HIV-specific antibody mutations by engineering B cell maturation. Science 366(2019).

15. Molinos-Albert, L.M., et al. Anti-V1/V3-glycan broadly HIV-1 neutralizing antibodies in a post-treatment controller. Cell Host Microbe 31, 1275–1287 e1278 (2023).

16. Sanders, R.W., et al. A next-generation cleaved, soluble HIV-1 Env trimer, BG505 SOSIP.664 gp140, expresses multiple epitopes for broadly neutralizing but not non-neutralizing antibodies. PLoS Pathog 9, e1003618 (2013).

17. Bonsignori, M., et al. Staged induction of HIV-1 glycan-dependent broadly neutralizing antibodies. Sci Transl Med 9(2017).

18. Walker, L.M., et al. Broad neutralization coverage of HIV by multiple highly potent antibodies. Nature 477, 466–470 (2011).

19. Barnes, C.O., et al. Structural characterization of a highly-potent V3-glycan broadly neutralizing antibody bound to natively-glycosylated HIV-1 envelope. Nat Commun 9, 1251 (2018).

20. Silva, M., et al. A particulate saponin/TLR agonist vaccine adjuvant alters lymph flow and modulates adaptive immunity. Sci Immunol 6, eabf1152 (2021).

21. Steichen, J.M., et al. HIV Vaccine Design to Target Germline Precursors of Glycan-Dependent Broadly Neutralizing Antibodies. Immunity 45, 483–496 (2016).

22. Sajadi, M.M., et al. A comprehensive engineering strategy improves potency and manufacturability of a near pan-neutralizing antibody against HIV. bioRxiv, 2024.2010.2014.618178 (2024).

23. Simonich, C.A., et al. Kappa chain maturation helps drive rapid development of an infant HIV-1 broadly neutralizing antibody lineage. Nat Commun 10, 2190 (2019).

24. MacLeod, D.T., et al. Early Antibody Lineage Diversification and Independent Limb Maturation Lead to Broad HIV-1 Neutralization Targeting the Env High-Mannose Patch. Immunity 44, 1215–1226 (2016).

25. Swanson, O., et al. Rapid selection of HIV envelopes that bind to neutralizing antibody B cell lineage members with functional improbable mutations. Cell Rep 36, 109561 (2021).

26. Kumar, S., et al. An HIV-1 Broadly Neutralizing Antibody from a Clade C-Infected Pediatric Elite Neutralizer Potently Neutralizes the Contemporaneous and Autologous Evolving Viruses. J Virol 93(2019).

27. Steichen, J.M., et al. Vaccine priming of rare HIV broadly neutralizing antibody precursors in nonhuman primates. Science 384, eadj8321 (2024).

28. Escolano, A., et al. Sequential immunization of macaques elicits heterologous neutralizing antibodies targeting the V3-glycan patch of HIV-1 Env. Sci Transl Med 13, eabk1533 (2021).

29. Cirelli, K.M., et al. Slow Delivery Immunization Enhances HIV Neutralizing Antibody and Germinal Center Responses via Modulation of Immunodominance. Cell 180, 206 (2020).

30. Moore, P.L., et al. Evolution of an HIV glycan-dependent broadly neutralizing antibody epitope through immune escape. Nat Med 18, 1688–1692 (2012).

31. Alam, S.M., et al. Mimicry of an HIV broadly neutralizing antibody epitope with a synthetic glycopeptide. Sci Transl Med 9(2017).

32. Henderson, R., et al. Structural basis for breadth development in the HIV-1 V3-glycan targeting DH270 antibody clonal lineage. Nat Commun 14, 2782 (2023).

33. Wang, H., et al. Asymmetric recognition of HIV-1 Envelope trimer by V1V2 loop-targeting antibodies. Elife 6(2017).

34. Zeng, W.F., Cao, W.Q., Liu, M.Q., He, S.M. & Yang, P.Y. Precise, fast and comprehensive analysis of intact glycopeptides and modified glycans with pGlyco3. Nat Methods 18, 1515–1523 (2021).

35. Bern, M., Kil, Y.J. & Becker, C. Byonic: advanced peptide and protein identification software. Curr Protoc Bioinformatics Chapter 13, 13 20 11–13 20 14 (2012).

36. Lee, J.H., et al. Long-primed germinal centres with enduring affinity maturation and clonal migration. Nature 609, 998–1004 (2022).

37. von Boehmer, L., et al. Sequencing and cloning of antigen-specific antibodies from mouse memory B cells. Nat Protoc 11, 1908–1923 (2016).

38. Viant, C., Escolano, A., Chen, S.T. & Nussenzweig, M.C. Sequencing, cloning, and antigen binding analysis of monoclonal antibodies isolated from single mouse B cells. STAR Protoc 2, 100389 (2021).

39. Williams, W.B., et al. Fab-dimerized glycan-reactive antibodies are a structural category of natural antibodies. Cell 184, 2955–2972 e2925 (2021).

40. Gaebler, C., et al. Evolution of antibody immunity to SARS-CoV-2. Nature 591, 639–644 (2021).

41. Diskin, R., Marcovecchio, P.M. & Bjorkman, P.J. Structure of a clade C HIV-1 gp120 bound to CD4 and CD4-induced antibody reveals anti-CD4 polyreactivity. Nat Struct Mol Biol 17, 608–613 (2010).

42. Sarzotti-Kelsoe, M., et al. Optimization and validation of the TZM-bl assay for standardized assessments of neutralizing antibodies against HIV-1. J Immunol Methods 409, 131–146 (2014).

43. Montefiori, D.C. Measuring HIV neutralization in a luciferase reporter gene assay. Methods Mol Biol 485, 395–405 (2009).

44. Scheres, S.H. RELION: implementation of a Bayesian approach to cryo-EM structure determination. J Struct Biol 180, 519–530 (2012).

45. Pettersen, E.F., et al. UCSF Chimera--a visualization system for exploratory research and analysis. J Comput Chem 25, 1605–1612 (2004).

46. Patel, A., Toso, D., Litvak, A. & Nogales, E. Efficient graphene oxide coating improves cryo-EM sample preparation and data collection from tilted grids. bioRxiv, 2021.2003.2008.434344 (2021).

47. Waterhouse, A., et al. SWISS-MODEL: homology modelling of protein structures and complexes. Nucleic Acids Res 46, W296–W303 (2018).

48. Meng, E.C., et al. UCSF ChimeraX: Tools for structure building and analysis. Protein Sci 32, e4792 (2023).

49. Emsley, P. & Cowtan, K. Coot: model-building tools for molecular graphics. Acta Crystallogr D Biol Crystallogr 60, 2126–2132 (2004).

50. Chen, V.B., et al. MolProbity: all-atom structure validation for macromolecular crystallography. Acta Crystallogr D Biol Crystallogr 66, 12–21 (2010).

51. Agirre, J., et al. Privateer: software for the conformational validation of carbohydrate structures. Nat Struct Mol Biol 22, 833–834 (2015).

52. Barad, B.A., et al. EMRinger: side chain-directed model and map validation for 3D cryo-electron microscopy. Nat Methods 12, 943–946 (2015).

