## Extended Data Figures for "Rapid elicitation of a new class of neutralizing N332-glycan independent V3-glycan antibodies against HIV-1 in nonhuman primates"

**a**

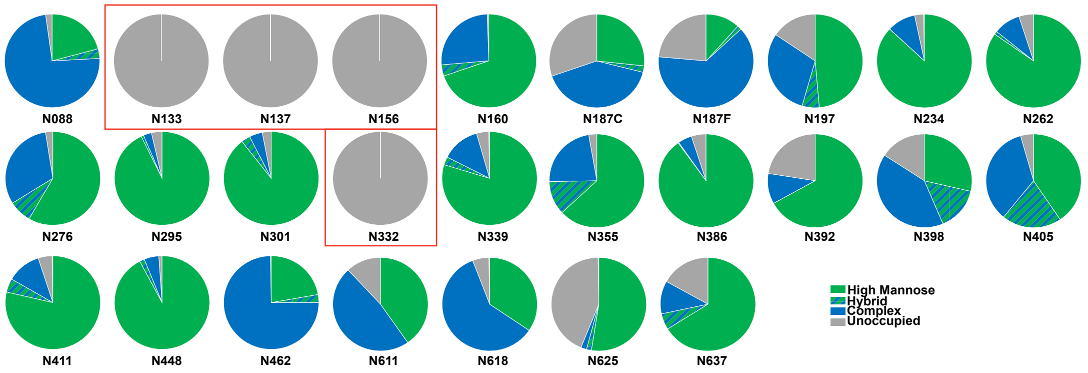
N088
N133
N137
N156
N160
N187C
N187F
N197
N234
N262
N276
N295
N301
N332
N339
N355
N386
N392
N398
N405
N411
N448
N462
N611
N618
N625
N637
High Mannose
Hybrid
Complex
Unoccupied

**b**

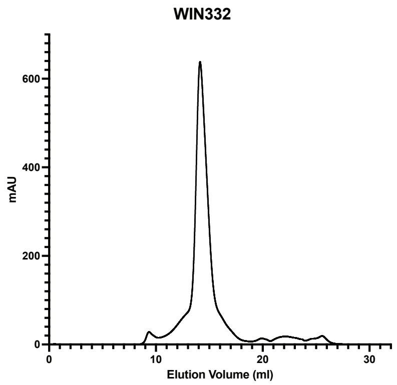
WIN332
mAU
Elution Volume (ml)

**c**

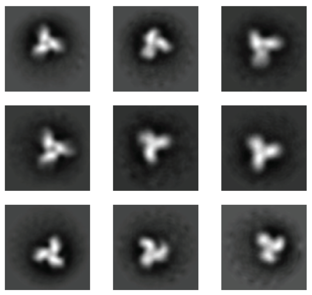

**d**

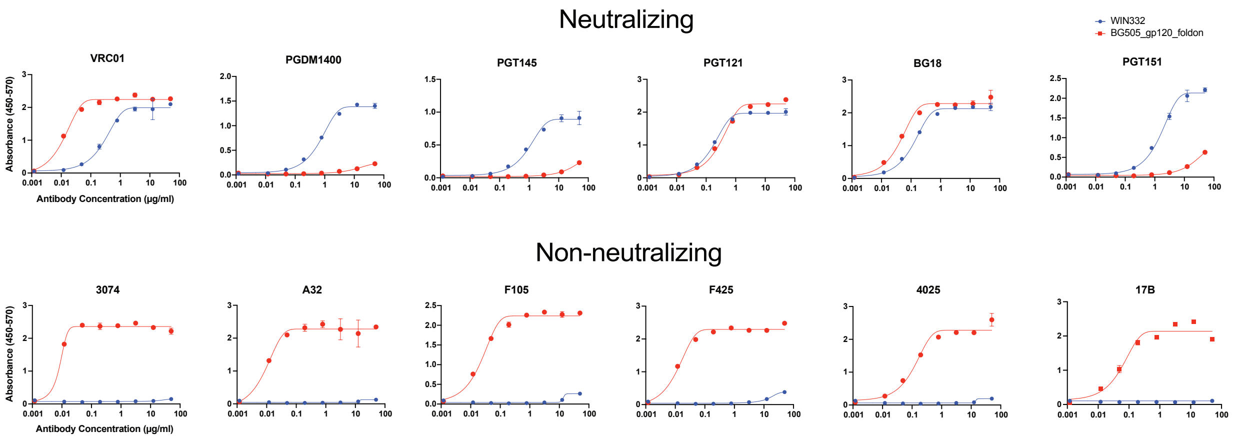
Neutralizing
WIN332
BG505_gp120_foldon
VRC01
PGDM1400
PGT145
PGT121
BG18
PGT151
Absorbance (450-570)
Antibody Concentration (µg/ml)
Non-neutralizing
3074
A32
F105
F425
4025
17B

**e**

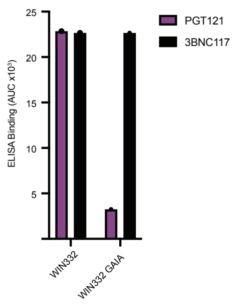
PGT121
3BNC117
ELISA Binding (AUC x10⁵)
WIN332
WIN332 GAIA

**Extended Data Figure 1: Biophysical and antigenic characterization of WIN332**

**a.** Mass spectrometry analysis of WIN332 glycosylation. Pie charts show the glycan composition of WIN332 Potential N-glycosylation sites (PNGS). Gray pie charts indicate unoccupied PNGSs. The frequencies of high mannose, hybrid and complex-type glycans found at each PNGS are shown in green, dashed green-blue, and blue respectively. **b.** Size exclusion chromatography (SEC) showing elution volume for fractions containing WIN332. **c.** Negative staining electron microscopy (nsEM) 2D classes of WIN332 showing the expected propeller shape. **d.** Antigenic profiling of WIN332 by ELISA. ELISA plots show the binding of different human bNAbs and the lack of binding of non-neutralizing antibodies to WIN332. **e**. ELISA controls for the experiments in Fig.1f.

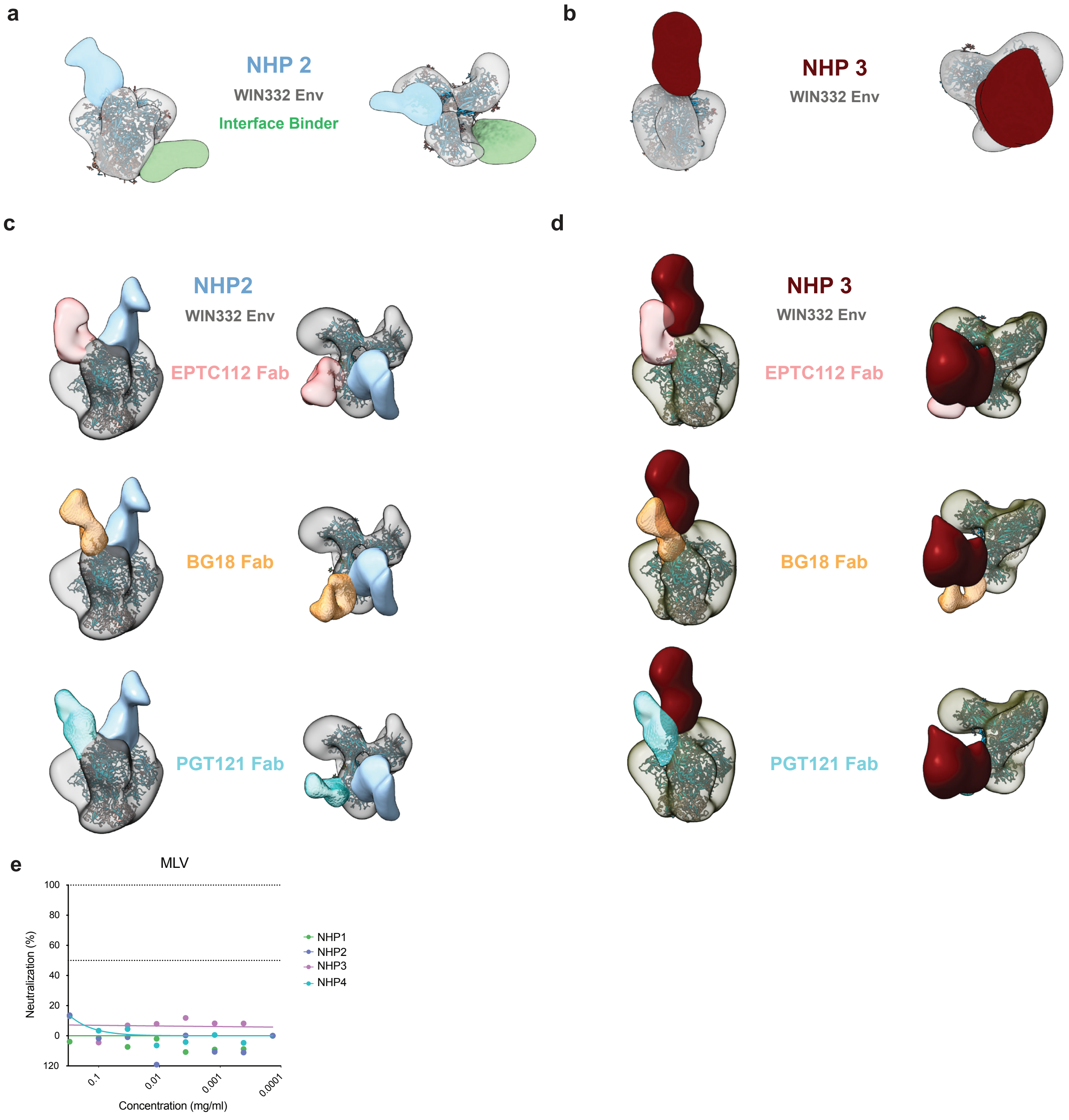


**Extended Data Figure 2: EMPEM analiysis of the serum from WIN332-primed NHPs.**

**a, b.** Negative staining electron microscopy (nsEM) densities of WIN332 Env (gray) bound to Fabs from NHP2 (blue and green) (a) and NHP3 (red) (b). WIN332 density is modelled on PDB6ORO. **c,d.** Comparison of the NHP2 (c) and NHP3 (d) top binder Fab densities with published Fabs EPTC112 (pink), BG18 (orange), and PGT121 (cyan). **e.** TZM-bl neutralization assay to determine the activity of the NHP serum in Fig. 2h against the control Murine Leukemia Virus (MLV).

a

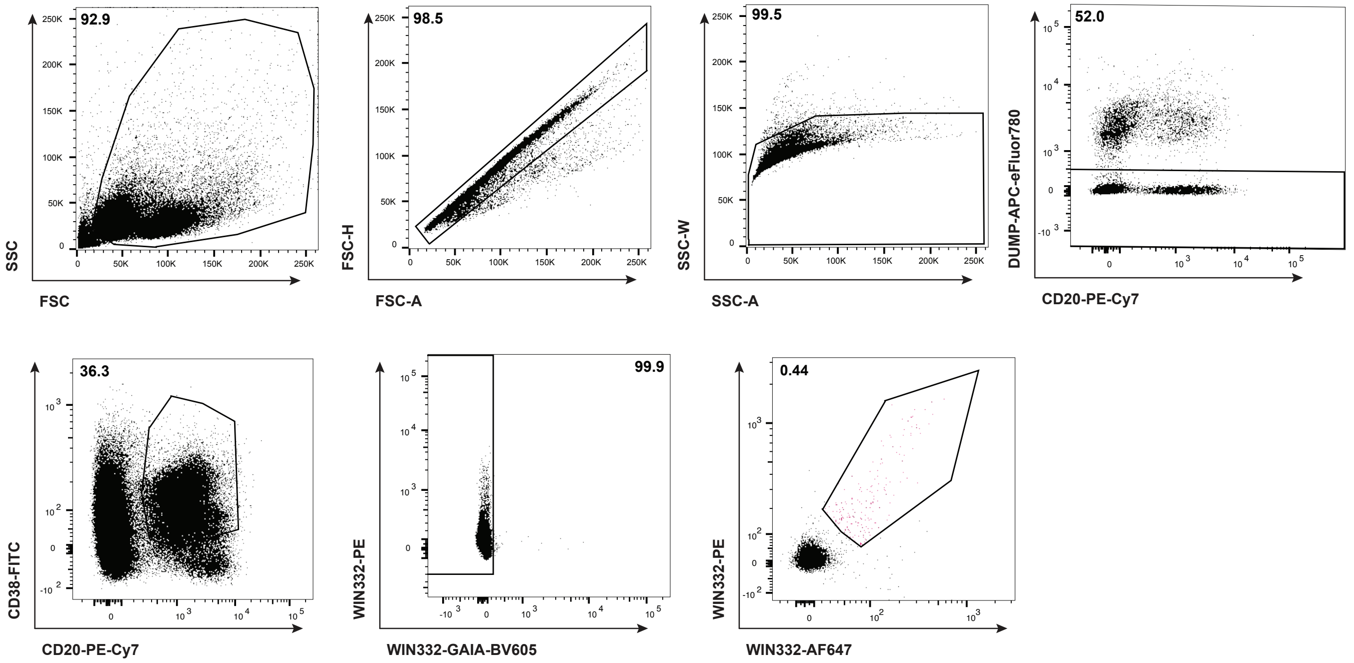


b

| HC | |
|---|---|
| bNAb | Motif |
| BG18 | FGVV |
| PGT121 | RxY |
| EPTC112 | SxW |
| PCDN76 | WSG |
| DH270.6 | YYxS |
| PGT128 | LxYxD/E |
| QA013.2 | DDR |

| LC | |
|---|---|
| bNAb | Motif |
| PGT121 | QxDSS |
| EPTC112 | SYxG |
| PCDN76 | YxSS/N |
| DH1030 | THL/I |
| AIMS-P01 | AWDxS |
| BF520 | YxN/NxY |

**Extended Data Figure 3: Gating strategy to isolate single WIN332-specific B cells from NHP LN biopsies.**

**a.** Representative flow cytometry plots showing the gating strategy used to isolate single WIN332-specific B cells from cell homogenates of WIN332-primed NHP LNs. A WIN332 protein with mutations at the GDIR motif (WIN332-GAIA) was used as a negative bait. **b.** Human bNAb motifs used to analyze NHP immunoglobulin sequences in Fig. 3e.

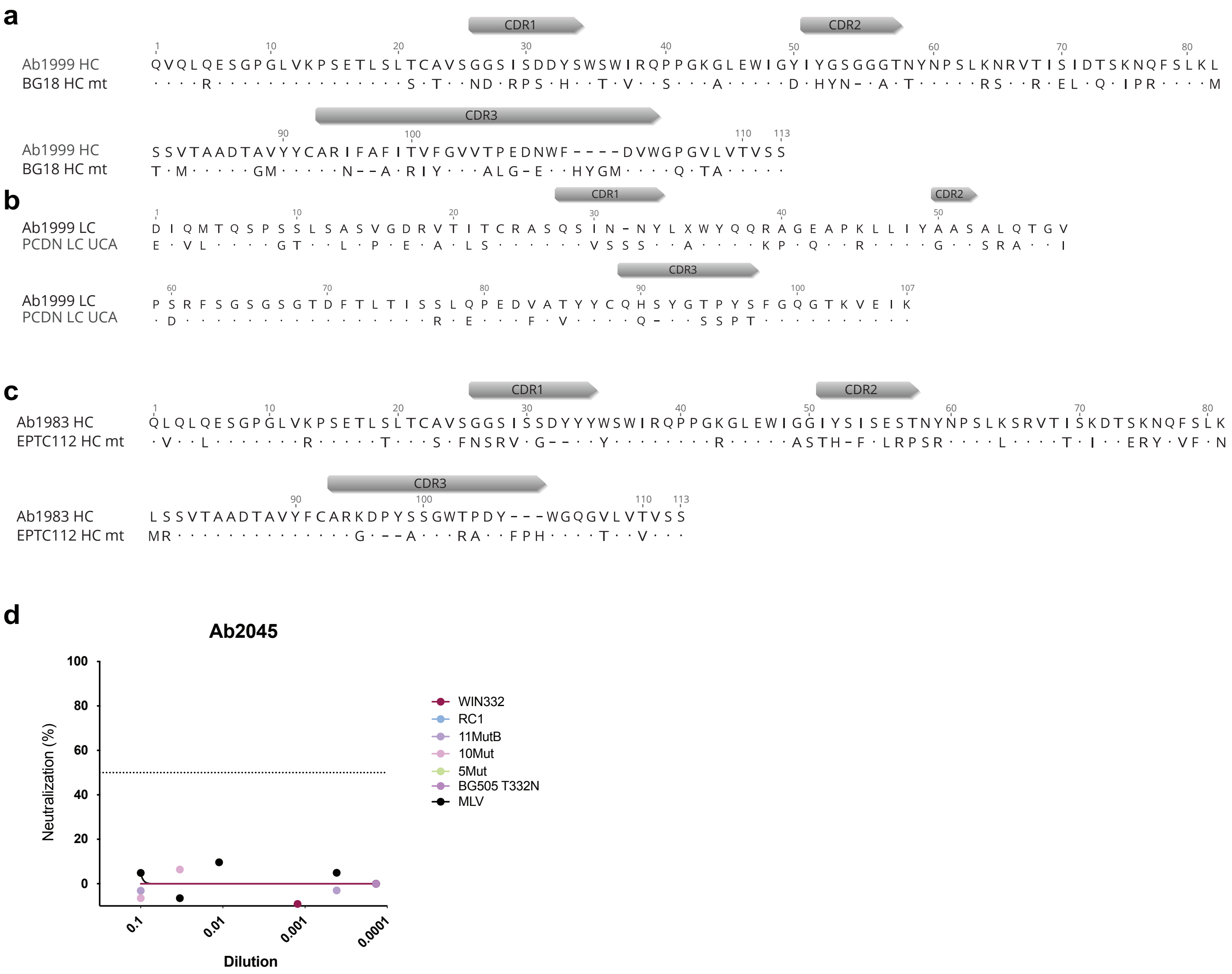


**Extended Data Figure 4: Sequence identity between Ab1999 and 1983 with human V3-glycan bNAbs.**

**a,b.** Amino acid sequence alignment of Ab1999 HC with BG18 HC mt (a) and of Ab1999 LC with PCDN.76 UCA LC (b). **c.** Amino acid sequence alignment of Ab1983 HC with EPTC112 iGL HC. Dots represent identical amino acids. Dashes represent absent amino acids. **c.** Control TZM-bl assay using a non-neutralizing NHP antibody isolated from one of the WIN332-primed NHPs.

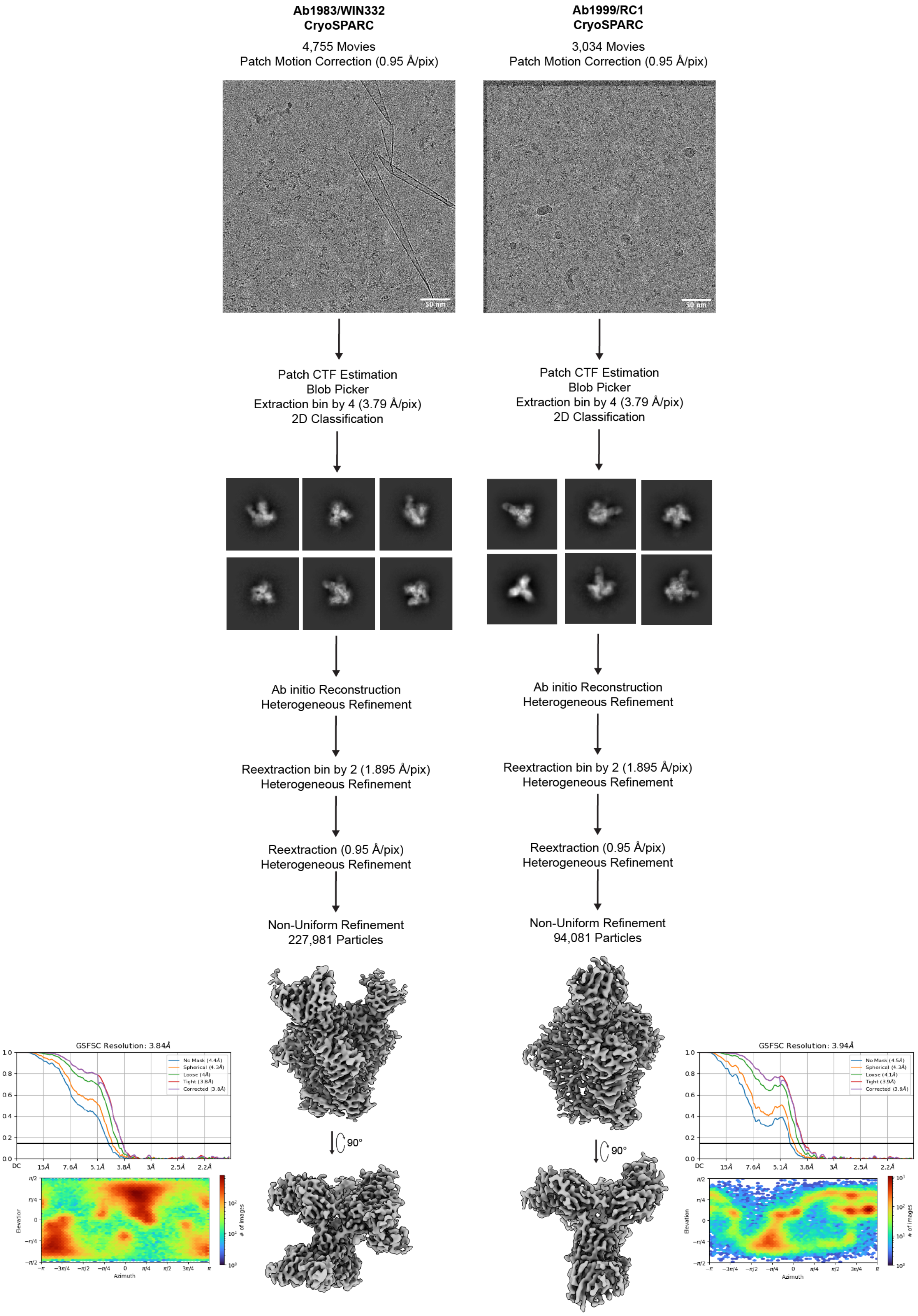


**Extended Data Figure 5: Cryo-EM data processing scheme for Ab1983/WIN332 and Ab1999/RC1 complexes.**

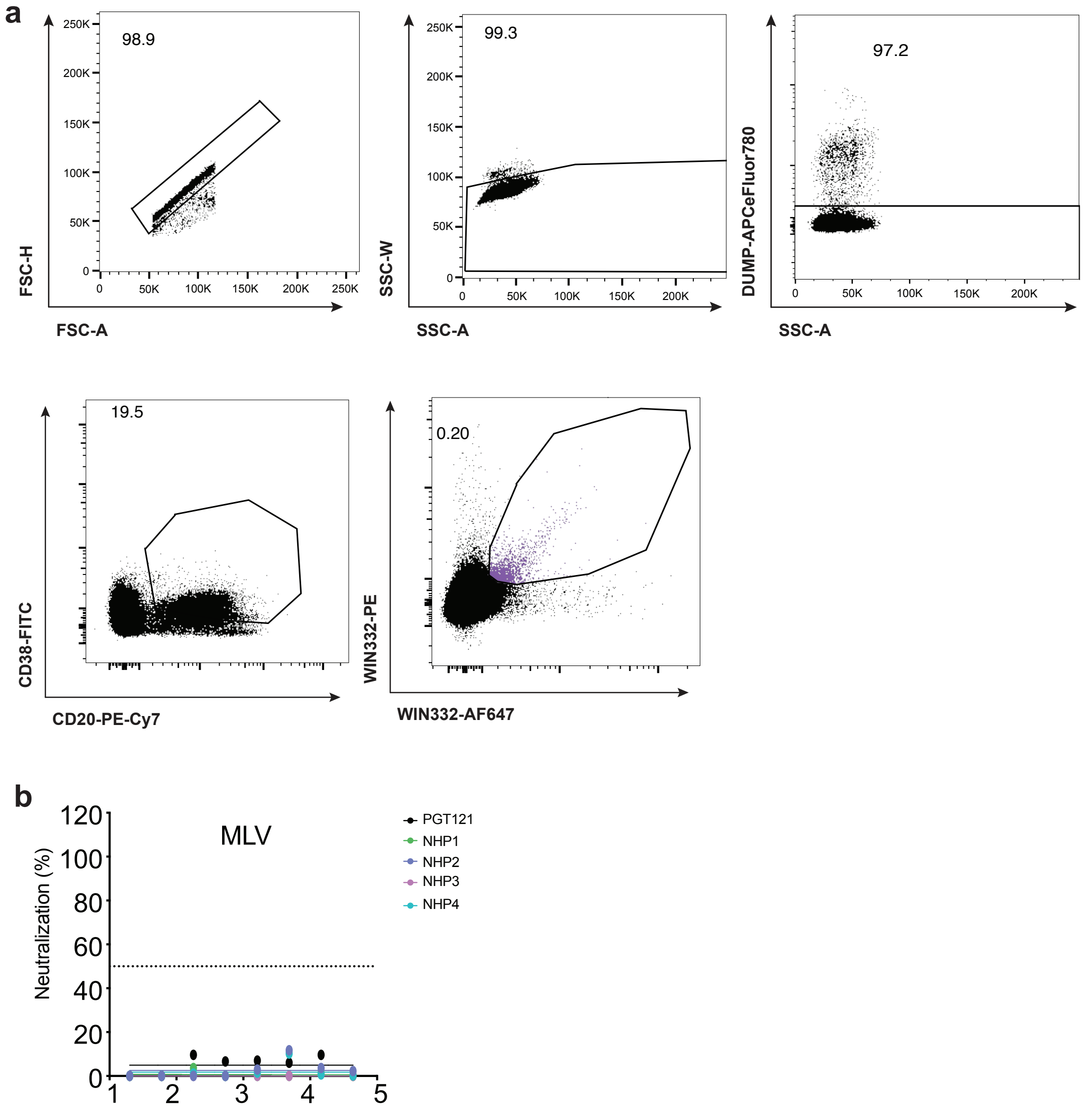


**Extended Data Figure 6: Gating strategy to isolate single WIN332-specific B cells from NHP LN biopsies after sequential immunization.**

**a.** Flow cytometry plots showing the gating strategy used to isolate single WIN332-specific B cells from a pool of cell homogenates prepared from LN biopsies from sequentially immunized NHPs. **b.** TZM-bl neutralization assay to determine the activity of the NHP serum in Fig. 6h against the control Murine Leukemia Virus (MLV).

| Protein | Deleted PNGS | Other modifications |
|---|---|---|
| WIN332 | 133, 137, 156, 332 | V134Y, T135A, N136P, I138L, T139L, D140S, D141N, N156A, T320F, Q328M, N332Q, T415V, MD39* |
| WIN332-AviBio | 133, 137, 156, 332 | V134Y, T135A, N136P, I138L, T139L, D140S, D141N, N156A, T320F, Q328M, N332Q, T415V, Avitag, MD39* |
| WIN332-GAIA-AviBio | 133, 137, 156, 332 | V134Y, T135A, N136P, I138L, T139L, D140S, D141N, N156A, T320F, Q328M, N332Q, T415V, GDIR/GAIA, Avitag, MD39* |
| WIN332-Glycan KO | 133, 137, 156, 301, 332 | V134Y, T135A, N136P, I138L, T139L, D140S, D141N, N156A, N301A, T320F, Q328M, N332Q, T415V, MD39* |
| RC1 | 133, 137, 156 | V134Y, T135A, N136P, I138L, T139L, D140S, D141N, N156A, T320F, Q328M, T415V, MD39* |
| 11MutB | 133, 137 | V134Y, T135A, N136P, I138L, T139L, D140S, D141N, T320F, Q328M, T415V, MD39* |
| 10Mut | 133, 137 | V134Y, T135A, N136P, N137F, I138L, T139I, D140N, T320F,Q328M, MD39* |
| 7Mut | 133, 137 | V134Y, T135A, N136P, I138L, T139I, D140N, MD39* |
| 5Mut | - | V134Y, N136P, I138L, D140N, MD39* |
| BG505 | - | MD39* |
| BG505 gp120 | - | MD39* |
| BG505-AviBio | - | Avitag, MD39* |
| CH119 | - | MD39* |
| Q23.17 | - | MD39* |
| AD8 | - | MD39* |
| CNE55 | - | MD39* |
| 001428 | - | MD39* |
| CE0217 | - | MD39* |
| BJOX002000 | - | MD39* |

**Extended Data Table 1: Env proteins used in the study and specific modifications.**

PNGS: Potential N-Glycosylation Site.

| | Heavy Chain | | | | | Light Chain | | | |
|---|---|---|---|---|---|---|---|---|---|
| Ab name | V Gene | D gene | J Gene | CDR sequence AA | CDR length (aa) | V Gene | J Gene | CDR sequence aa | CDR length (aa) |
| 1951 | IGHV5-43*02 | IGHD6-13*01 | IGHJ1*01 | AKGYSRWSEYFEF | 13 | IGLV1-85*01 | IGLJ6*01 | QSYDSSLSADV | 11 |
| 1952 | IGHV5-20*02 | IGHD6-13*01 | IGHJ1*01 | ATQYSSWSLPEYFEE | 15 | IGLV1-65*01 | IGLJ3*01 | SAWDNSLSAQL | 11 |
| 1953 | IGHV3-124*02 | IGHD5-36*01 | IGHJ4*01 | ARDGGGYSYGPIDY | 14 | IGLV1-85*01 | IGLJ6*01 | QSYDSSLSAHV | 11 |
| 1954 | IGHV4-106*01 | IGHD3-22*02 | IGHJ5-2*02 | ARDRGTPWGSSLDV | 14 | IGLV1-85*01 | IGLJ6*01 | QSYDSSLSAHV | 11 |
| 1955 | IGHV5-43*02 | IGHD6-31*01 | IGHJ5-1*02 | AKGDSGWFWDTWLDV | 15 | IGLV1-85*01 | IGLJ6*01 | QSYDSSLTAHV | 11 |
| 1956 | IGHV4-173*01 | IGHD6-25*01 | IGHJ4*01 | ASLLGSWNWDDIDY | 14 | IGLV1-85*01 | IGLJ2*01 | QSYDNSLSAHL | 11 |
| 1957 | IGHV4-106*01 | IGHD1-20*01 | IGHJ1*01 | ARDLAGTPWGLYLEL | 15 | IGLV1-85*01 | IGLJ2*01 | QSYDSSLSAQV | 11 |
| 1958 | IGHV4-106*01 | IGHD3-22*02 | IGHJ5-2*02 | ARGWGDYFGTYNSLDV | 16 | IGLV1-85*01 | IGLJ2*01 | QSYDSSLSVL | 10 |
| 1959 | IGHV1S2*01 | IGHD3-3*02 | IGHJ4*01 | ARGVGGPWSYFDY | 13 | IGLV1-85*01 | IGLJ6*01 | QSYDNSLSADV | 11 |
| 1960 | IGHV1-200*01 | IGHD3-28*01 | IGHJ6*01 | ARMSGYYTDGLDS | 13 | IGLV1-85*01 | IGLJ2*01 | QSYDNSLSVQV | 11 |
| 1961 | IGHV7-114*03 | IGHD1-32*01 | IGHJ4*01 | ASYRNLDY | 8 | IGLV3-40*01 | IGLJ3*01 | QVWDSSSDQVL | 11 |
| 1962 | IGHV3-116*02 | IGHD3-3*01 | IGHJ5-1*02 | TRVRVLQFLEDVRSHWWFDV | 20 | IGLV1-85*01 | IGLJ6*01 | QSYDSSLSAHV | 11 |
| 1975 | IGHV5-43*02 | IGHD6-13*01 | IGHJ4*01 | AKGYSRWLEYFDY | 13 | IGLV1-85*01 | IGLJ6*01 | QSYDSSLTADV | 11 |
| 1976 | IGHV3-115*02 | IGHD6-25*01 | IGHJ4*01 | ARHSIGGGGSWNYFDY | 16 | IGLV11-117*01 | IGLJ3*01 | QVYDSSAVL | 9 |
| 1977 | IGHV1-180*01 | IGHD4-23*01 | IGHJ1*01 | TRGHYEYSNYEYFEF | 15 | IGLV3-40*01 | IGLJ6*01 | QVWDSSSDDV | 10 |
| 1978 | IGHV3-110*02 | IGHD6-31*01 | IGHJ5-2*02 | ARHSSGWLSDSLDV | 14 | IGLV1-85*01 | IGLJ2*01 | QSYDSSLRALL | 11 |
| 1979 | IGHV3-110*02 | IGHD6-13*01 | IGHJ5-2*02 | ARHSSGWLSDSLDV | 14 | IGLV3-40*01 | IGLJ3*01 | QVWDNNSDHYLL | 12 |
| 1981 | IGHV3-184*01 | IGHD6-31*01 | IGHJ4*01 | SRVGSGWYPYFDY | 13 | IGLV3-40*01 | IGLJ2A*01 | QVWDSSSDHPV | 11 |
| 1982 | IGHV7-114*03 | IGHD6-13*01 | IGHJ4*01 | VRRGYSSWSDFDY | 13 | IGLV3-40*01 | IGLJ3*01 | QVWDSSSDHPL | 11 |
| 1983 | IGHV4S13*01 | IGHD6-31*01 | IGHJ4*01 | ARKDPYSSGWTPDY | 14 | IGLV9-84*01 | IGLJ2A*01 | GADHGTGSSFVWV | 13 |
| 1998 | IGHV3-8*01 | IGHD6-25*01 | IGHJ4*01 | AKGDSYSGSSHFDY | 14 | IGKV2S16*01 | IGKJ3*01 | MQGIEYFT | 8 |
| 1999 | IGHV4-106*01 | IGHD3-3*01 | IGHJ5-1*02 | ARIFAFITVFGVVTPEDNWFDV | 22 | IGKV1-74*01 | IGKJ2*01 | QHSYGTPYS | 9 |
| 2000 | IGHV3-178*05 | IGHD6-31*01 | IGHJ4*01 | ARKRSGGWYQYRGDY | 15 | IGKV2-86*01 | IGKJ4*01 | MQYTHIPLT | 9 |
| 2001 | IGHV3-124*02 | IGHD4-23*01 | IGHJ4*01 | ARPRMNTVTWRFDY | 14 | IGKV2-104*02 | IGKJ3*01 | MQGLEFFT | 8 |
| 2003 | IGHV1-198*02 | IGHD6-25*01 | IGHJ5-2*02 | ASDTYSDSLIGDV | 13 | IGKV2-65*01 | IGKJ3*01 | GQGTHLPFT | 9 |
| 2004 | IGHV3-103*04 | IGHD5-12*01 | IGHJ4*01 | AKVVSLGYSYSPTFDY | 16 | IGKV2-104*02 | IGKJ1*01 | MQGLELWT | 8 |
| 2005 | IGHV3-183*01 | IGHD1-1-1*01 | IGHJ5-1*02 | SRVGDLFRYSWNNWFDV | 17 | IGKV1-32*05 | IGKJ1*01 | QQYNSLPPWT | 10 |
| 2006 | IGHV5-20*02 | IGHD5-24*01 | IGHJ6*01 | AKPTYSGYKGWGNGLDS | 17 | IGKV1-94*01 | IGKJ2*01 | QQDYTTPYS | 9 |
| 2007 | IGHV1-111*02 | IGHD6-31*01 | IGHJ5-2*02 | TTMWTPGFTNGWYNSLDV | 18 | IGKV1-32*05 | IGKJ2*01 | QQYNSLPFS | 9 |
| 2008 | IGHV3-182*05 | IGHD3-28*02 | IGHJ4*01 | ARHEHYFGSSSHFAY | 15 | IGKV1-22*01 | IGKJ3*01 | LQYISSPFT | 9 |
| 2044 | (h)IGHV4-4*02 | (h)IGHD4-17*01 | (h)IGHJ5*02 | ARHPPWDYEDDYGYYRGGRFDV | 22 | IGLV1-85*01 | IGLJ6*01 | QSYDSSLSADV | 11 |
| 2045 | IGHV3-110*01 | IGHD2-33*01 | IGHJ4*01 | ARQDCGDSDCSSSFDY | 16 | IGLV3-36*01 | IGLJ3*01 | QVWDSSSDHVL | 11 |
| 2053 | IGHV3-182*05 | IGHD2-39*01 | IGHJ4*01 | AKESRYCSRGVCPEFDY | 17 | IGLV4-97*01 | IGLJ6*01 | QTWTTGIHV | 9 |
| 2054 | IGHV1-111*01 | IGHD3-28*01 | IGHJ4*01 | ATEGDTSRSRPTYYYDTGYFT | 21 | IGLV8-125*01 | IGLJ6*01 | MLYMGSGISV | 10 |
| 2055 | IGHV3-178*01 | IGHD2-21*01 | IGHJ4*01 | ARDTREYCPGSGCRYFDY | 18 | IGLV1-64*01 | IGLJ2*01 | QSYDSRLSVGL | 11 |
| 2056 | IGHV4-80*03 | IGHD4-29*01 | IGHJ1*01 | ARGDYGSIRGYFEF | 14 | IGLV3-36*02 | IGLJ1*01 | QVWDSSSDHYI | 11 |
| 2057 | IGHV1-151*01 | IGHD3-16*01 | IGHJ4*01 | ARGARGDYSGSYYYTHFDY | 19 | IGLV2-19*03 | IGLJ1*01 | SSYADSNTFI | 10 |
| 2087 | IGHV3S4*01 | IGHD2-8*01 | IGHJ6*01 | TRDRDCSGGVCYAGSEYYYGLDS | 23 | IGLV3-36*02 | IGLJ2A*01 | QVWDSSSDHWV | 11 |
| 2088 | IGHV3-178*01 | IGHD4-29*01 | IGHJ4*01 | ARDWVAATSRAVYYFDY | 17 | IGLV8-125*01 | IGLJ3*01 | MLYMGSGISL | 10 |
| 2089 | IGHV3-182*05 | IGHD2-8*01 | IGHJ5-1*02 | AKDPLGYCSGGSCYSGSGWFDP | 22 | IGLV9-84*01 | IGLJ1*01 | GADHGTGSSFVYI | 13 |
| 2090 | IGHV3-110*02 | IGHD2-15*01 | IGHJ4*01 | ARQYCSSTYCSSGFDY | 16 | IGLV3-36*01 | IGLJ3*01 | QVWDSSSDHPL | 11 |
| 2092 | IGHV3S4*01 | IGHD2-8*01 | IGHJ6*01 | ARDRDCSGGVCYAGSDYYYGLDS | 23 | IGLV3-36*02 | IGLJ2A*01 | QVWDSSSDHWV | 11 |
| 2094 | IGHV4-80*04 | IGHD4-29*01 | IGHJ1*01 | ARGDYGAIRGYFEF | 14 | IGLV3-36*02 | IGLJ1*01 | QVWDSSSDHYI | 11 |
| 2097 | IGHV3-136*01 | IGHD2-8*01 | IGHJ4*01 | TRDRDCTGGVCYAPRQTYYYDSGYDY | 26 | IGLV3-36*02 | IGLJ1*01 | QVWDSSSDHYI | 11 |
| 2187 | IGHV4-106*01 | IGHD3-3*01 | IGHJ4*01 | AGQLYNFWSAYQSNFDY | 17 | (h)IGKV1-9*01 | (h)IGKJ4*01 | QQRNSYPLT | 9 |
| 2205 | IGHV3-116*02 | IGHD5-24*01 | IGHJ5-1*02 | ARDRYSAYSPENWFDV | 16 | IGKV1-22*01 | IGKJ1*01 | QQYSSSPRT | 9 |
| 2207 | IGHV1-198*03 | IGHD3-16*01 | IGHJ5-1*01 | ARGRWGGSHHSRPYFYGLEY | 20 | IGKV2-65*01 | IGKJ4*01 | GQGTHLFT | 8 |
| 2209 | IGHV1-156*01 | IGHD6-31*01 | IGHJ1*01 | ARGPRSSGWYHEYFDF | 16 | IGLV2S9*01 | IGLJ1*01 | CSYRSGSTYI | 10 |
| 2210 | IGHV3S4*01 | IGHD2-8*01 | IGHJ6*01 | TRDRDCSGGVCYAGSEYYYGLDS | 23 | IGLV3-36*02 | IGLJ2A*01 | QVWDSSSDHWV | 11 |
| 2211 | IGHV3-136*01 | IGHD2-8*01 | IGHJ4*01 | TRDRDCTGGVCYAPRQTFYYDSGFDY | 26 | IGLV3-36*02 | IGLJ1*01 | QVWDSSSDHYI | 11 |
| 2212 | IGHV3-178*02 | IGHD2-21*02 | IGHJ5-1*02 | AKDMRCTGSGCYGNWFDV | 18 | !"#$%&''()* | !"#+%()* | QSADSSGNHVL | 11 |
| 2273 | IGHV5-10-1*04 | IGHD4-23*01 | IGHJ1*01 | ARGDYGGIREFFEF | 14 | IGLV3-21*01 | IGLJ2*01 | QVWDTSSAHPV | 11 |
| 2315 | IGHV4-4*02 | IGHD6-19*01 | IGHJ5*02 | ARDPGGGWTYNRFDV | 15 | IGLV3-21*02 | IGLJ2*01 | QVWDSGSKYVL | 11 |
| 2316 | IGHV3-71*01 | IGHD6-13*01 | IGHJ4*02 | ATSLGGAAAGTFLY | 14 | IGLV5-48*02 | IGLJ2*01 | MIWHDIDFVL | 10 |
| 2346 | IGHV7-114*04 | IGHD4-29*02 | IGHJ4*01 | ASGRNYGDS | 9 | IGLV3-34*01 | IGLJ2*01 | QVWDSSSNL | 9 |
| 2348 | IGHV5-20*02 | IGHD3-28*02 | IGHJ4*01 | AMGLYGSGYPFYADY | 15 | IGLV1-65*01 | IGLJ2A*01 | SAWDNSLSAWV | 11 |
| 2349 | IGHV5-20*02 | IGHD3-3*02 | IGHJ5-2*01 | AKSEGAFWRGYYTGLLEV | 18 | IGLV1-60*01 | IGLJ1*01 | AAWDDSLSGYI | 11 |

**Extended Data Table 2: Features of monoclonal antibodies isolated from the WIN332-immunized macaques.**

| Heavy chain | | | |
|---|---|---|---|
| | | Primer name | Primer sequence |
| **1st PCR** | Forward | p1350 | ACAGGTGCCCACTCCCAGGTGCAG |
| | | p1351 | AAGGTGTCCAGTGTGARGTGCAG |
| | | p1352 | CCCAGATGGGTCCTGTCCCAGGTGCAG |
| | | p1353 | CAAGGAGTCTGTTCCGAGGTGCAG |
| | | VH5 LEADER-A | TTCTCCAAGGAGTCTGT |
| | | VH3 LEADER-A | TAAAAGGTGTCCAGTGT |
| | | VH3 LEADER-B | TAAGAGGTGTCCAGTGT |
| | | VH3 LEADER-C | TAGAAGGTGTCCAGTGT |
| | | VH4 LEADER-D | ATGAAACATCTGTGGTTCTT |
| | | VH3 LEADER-E | TACAAGGTGTCCAGTGT |
| | | VH3 LEADER-F | TTAAAGCTGTCCAGTGT |
| | Reverse | 3'Sall.JH1/4/5 | *GCTGAGGAGACGGTGACCAG* |
| | | 3'Sall.JH2 | *GCTGAGGAGATGGTGATTGGG* |
| | | 3'Sall.JH3 | *GCTGAAGAGACGGTGACCCTG* |
| | | 3'Sall.JH6 | *GCTGAGGAGACGGTGACGACG* |
| **2nd PCR** | Forward | p1355 | CTAGTAGCAACTGCAACCGGTGTACATTCCCAGGTGCAGCTGGTGCAG |
| | | p1356 | CTAGTAGCAACTGCAACCGGTGTACATTCCGAGGTGCAGCTGGTGCAG |
| | | p1357 | CTAGTAGCAACTGCAACCGGTGTACATTCCCAGGTTCAGCTGGTGCAG |
| | | p1358 | CTAGTAGCAACTGCAACCGGTGTACATTCCCAGGTCCAGCTGGTACAG |
| | | p1359 | CTAGTAGCAACTGCAACCGGTGTACATTCTGAGGTGCAGCTGGTGGAG |
| | | p1360 | CTAGTAGCAACTGCAACCGGTGTACATTCTCAGGTGCAGCTGGTGGAG |
| | | p1361 | CTAGTAGCAACTGCAACCGGTGTACATTCTGAGGTGCAGCTGTTGGAG |
| | | p1362 | CTAGTAGCAACTGCAACCGGTGTACATTCTCAGGTGCAGCTGGTGGAG |
| | | p1363 | CTAGTAGCAACTGCAACCGGTGTACATTCTGAAGTGCAGCTGGTGGAG |
| | | p1364 | CTAGTAGCAACTGCAACCGGTGTACATTCCCAGGTGCAGCTGCAGGAG |
| | | p1365 | CTAGTAGCAACTGCAACCGGTGTACATTCCCAGGTGCAGCTACAGCAGTG |
| | | p1366 | CTAGTAGCAACTGCAACCGGTGTACATTCCCAGCTGCAGCTGCAGGAG |
| | | p1367 | CTAGTAGCAACTGCAACCGGTGTACATTCCCAGGTACAGCTGCAGCAG |
| | Reverse | p1371 | CCGATGGGCCCTTGGTCGACGCTGAAGAGACGGTGACCATTG |
| | | p1372 | CCGATGGGCCCTTGGTCGACGCTGAGGAGACGGTGACCAG |
| | | p1373 | CCGATGGGCCCTTGGTCGACGCTGAGGAGACGGTGACCGTG |
| **Sequencing** | | RM_FWD_T4_Seq | GTAGCAACTGCAACCGGTGT |
| **Colony PCI** | Forward | Ab-sense | GCTTCGTTAGAACGCGGCTAC |
| | Reverse | p1354 | GGAAGGTGTGCACGCCGCTGGTC |
| | Sequencing | Ab-sense | GCTTCGTTAGAACGCGGCTAC |

Heavy Chain rescue primers 1st PCR FWD

| | | |
|---|---|---|
| G1 | VH1 LEADER-A | ATGGACTGGACCTGGAGGAT |
| | VH1 LEADER-B | ATGGACTGGACCTGGAGCAT |
| | VH1 LEADER-C | ATGGACTGGACCTGGAGAAT |
| | VH1 LEADER-D | GGTTCCTCTTTGTGGTTGGC |
| | VH1 LEADER-E | ATGGACTGGACCTGGAGGGT |
| | VH1 LEADER-F | ATGGACTGGATTTGGAGGAT |
| | VH1 LEADER-G | AGGTTCCTCTTTGTGGTGGCAG |
| G2 | VH3 LEADER-D | GCTATTTTTAAAGGTGTCCAGTGT |
| | VH4 LEADER-A | ATGAAACACCTGTGGTTCTTCC |
| | VH4 LEADER-B | ATGAAACACCTGTGGTTCTT |
| | VH4 LEADER-C | ATGAAGCACCTGTGGTTCTT |
| | VH5 LEADER-B | CCTCCACAGTGAGAGTCTG |
| | VH6 LEADER-A | ATGTCTGTCTCCTTCCTCATC |
| | VH7 LEADER-A | GGCAGCAGCAACAGGTGCCCA |

| Light chain (Igl) | | | |
|---|---|---|---|
| | | Primer name | Primer sequence |
| **1st PCR** | Forward | p1394 | GGTCCTGGGCCCAGTCTGTGCTG |
| | | p1395 | GGTCCTGGGCCCAGTCTGCCCTG |
| | | p1396 | GCTCTGTGACCTCCTATGAGCTG |
| | | p1397 | GGTCTCTCTCSCAGCYTGTGCTG |
| | | p1398 | GTTCTTGGGCCAATTTTATGCTG |
| | | p1399 | GGTCCAATTCYCAGGCTGTGGTG |
| | | p1400 | GAGTGGATTCTCAGACTGTGGTG |
| | Reverse | p1401 | CACCAGTGTGGCCTTGTTGGCTTG |
| **2nd PCR** | Forward | p1402 | CTAGTAGCAACTGCAACCGGTTCCTGGGCCCAGTCTGTGCTGACKCAG |
| | | p1403 | CTAGTAGCAACTGCAACCGGTTCCTGGGCCCAGTCTGCCCTGACTCAG |
| | | p1404 | CTAGTAGCAACTGCAACCGGTTCTGTGACCTCCTATGAGCTGACWCAG |
| | | p1405 | CTAGTAGCAACTGCAACCGGTTCTCTCTCSCAGCYTGTGCTGACTCA |
| | | p1406 | CTAGTAGCAACTGCAACCGGTTCTTGGGCCAATTTTATGCTGACTCAG |
| | | p1407 | CTAGTAGCAACTGCAACCGGTTCCAATTCYCAGRCTGTGGTGACYCAG |
| | Reverse | p1409 | GGCTTGAAGCTCCTCACTCGAGGGYGGGAACAGAGTG |
| **Sequencing** | | p1409 | GGCTTGAAGCTCCTCACTCGAGGGYGGGAACAGAGTG |
| **Colony PCR** | Forward | Ab-sense | GCTTCGTTAGAACGCGGCTAC |
| | Reverse | p1409 | GGCTTGAAGCTCCTCACTCGAGGGYGGGAACAGAGTG |
| | Sequencing | Ab-sense | GCTTCGTTAGAACGCGGCTAC |

| Light chain (IgK) | | | |
|---|---|---|---|
| | | Primer name | Primer sequence |
| 1st PCR | Forward | p1374 | ATGAGGSTCCCYGCTCAGCTGCTGG |
| | | p1375 | CTCTTCCTCCTGCTACTCTGGCTCCCAG |
| | | p1376 | ATTTCTCTGTTGCTCTGGATCTCTG |
| | | p1377 | ATGACCCAGWCTCCABYCWCCCTG |
| | Reverse | p1378 | GTTTCTCGTAGTCTGCTTTGCTCA |
| 2nd PCR | Forward | p1379 | GTAGCAACTGCAACCGGTGTACATTCTGACATCCAGATGACCCAGTC |
| | | p1380 | GTAGCAACTGCAACCGGTGTACATTCAGACATCCAGTTGACCCAGTCT |
| | | p1381 | GTAGCAACTGCAACCGGTGTACATTGTGCCATCCGGATGACCCAGTC |
| | | p1382 | GTAGCAACTGCAACCGGTGTACATGGGGATATTGTGATGACCCAGAC |
| | | p1383 | GTAGCAACTGCAACCGGTGTACATGGGGATATTGTGATGACTCAGTC |
| | | p1384 | GTAGCAACTGCAACCGGTGTACATGGGGATGTTGTGATGACTCAGTC |
| | | p1385 | GTAGCAACTGCAACCGGTGTACATTCAGAAATTGTGTTGACACAGTC |
| | | p1386 | GTAGCAACTGCAACCGGTGTACATTCAGAAATAGTGATGACGCAGTC |
| | | p1387 | GTAGCAACTGCAACCGGTGTACATTCAGAAATTGTGTTGACGCAGTCT |
| | | p1388 | GTAGCAACTGCAACCGGTGTACATTCGGACATCGTGATGACCCAGTC |
| | Reverse | p1390 | GAAGACAGATGGTGCAGCCACCGTACGTTTGATYTCCACCTTGGTC |
| | | p1391 | GAAGACAGATGGTGCAGCCACCGTACGTTTGATCTCCAGCTTGGTC |
| | | p1392 | GAAGACAGATGGTGCAGCCACCGTACGTTTGATATCCACTTTGGTC |
| | | p1393 | GAAGACAGATGGTGCAGCCACCGTACGTTTAATCTCCAGTCGTGTC |
| **Sequencing** | | RM_FWD_T4_Seq | GTAGCAACTGCAACCGGTGT |
| **Colony PCR** | Forward | Ab-sense | GCTTCGTTAGAACGCGGCTAC |
| | Reverse | p1378 | GTTTCTCGTAGTCTGCTTTGCTCA |
| | Sequencing | Ab-sense | GCTTCGTTAGAACGCGGCTAC |

**Extended Data Table 3: Primers used to amplify rhesus macaque immunoglobulin genes from single Env-specific B cells.**

1st PCR

| Temperature | Time | |
|---|---|---|
| 95 | 30s | |
| 95 | 30s | 50 cycles |
| 46 | 30s | |
| 68 | 40s | |
| 68 | 5 min | |
| 10 | ∞ | |

2nd PCR

| Temperature | Time | |
|---|---|---|
| 95 | 30s | |
| 95 | 30s | 50 cycles |
| 55 | 30s | |
| 68 | 40s | |
| 68 | 5 min | |
| 10 | ∞ | |

**Extended Data Table 4: PCR protocols used to amplify rhesus macaque immunoglobulin genes from single Env-specific B cells.**

| Data Collection and Processing | RC1/Ab1999 | WIN332/Ab1983 |
|---|---|---|
| Electron Microscope | ThermoFisher Glacios | |
| Electron Detector | Falcon 4 | |
| Magnification | 150,000 | |
| Voltage (kV) | 200 | |
| Total Electron Exposure ($e^-/A^2$) | 50 | 40 |
| Defocus Range (µm) | 0.8 - 2.2 | |
| Pixel Size (Å) | 0.95 | |
| Symmetry | C1 | C1 |
| Number of final particle images | 94,082 | 227,981 |
| Map Resolution (Å) | 3.94 | 3.84 |
| FSC Threshold | 0.143 | 0.143 |
| Sharpening Factor | 99.6 | 136.7 |
| Model Building and Refinement | | |
| Initial models used | PDB:6ORO | PDB:6ORO |
| Model Composition | | |
| Protein Chains | 8 | 10 |
| Protein Residues | 2161 | 2155 |
| Ligands | 69 | 65 |
| Deviations from ideal (RMSD) | | |
| Bond lengths outlier (#) | 0.011 (0) | 0.010 (0) |
| Bond angles outlier (#) | 1.119 (21) | 1.110 (17) |
| Ramachandran Plot | | |
| Favored (%) | 99.14 | 99.43 |
| Allowed (%) | 0.86 | 0.48 |
| Outlier (%) | 0 | 0 |
| Validation | | |
| Molprobity score | 0.76 | 0.71 |
| Clashscore | 0.85 | 0.66 |
| Poor rotamer (%) | 0.35 | 0.16 |
| EMRinger score | 2.21 | 1.92 |

**Extended Data Table 5: Model statistics of Ab1983/WIN332 and Ab1999/RC1 structures.**
